# Genetic diversity and thermal performance in invasive and native populations of African fig flies

**DOI:** 10.1101/800938

**Authors:** Aaron A. Comeault, Jeremy Wang, Silas Tittes, Kristin Isbell, Spencer Ingley, Allen H. Hurlbert, Daniel R. Matute

**Author notes:** **Data availability:** genome assemblies and sequence data will be made available at appropriate NCBI archives and phenotypic data at Dryad upon acceptance.

## Abstract

During biological invasions, alien populations can suffer losses of genetic diversity that are predicted to negatively impact their fitness/performance. Using genome sequences, we show that invasive populations of the African fig fly, *Zaprionus indianus*, have lower levels of genetic diversity compared to native populations, and that genetic diversity is lost more in regions of the genome with low recombination rates. However, genetic diversity remains as high or higher in invasive populations than in populations of non-invasive congeneric species. We then use parameter estimates from thermal performance curves measured for 13 species of *Zaprionus* to show that *Z. indianus* has the broadest thermal niche of measured species, and that performance does not differ between invasive and native populations. These results illustrate how aspects of genetic diversity in invasive species can be decoupled from measures of fitness, and that a broad thermal niche may have helped facilitate *Z. indianus’s* range expansion.

## Introduction

Populations of invasive species can experience extreme demographic histories. For example, processes such as bottlenecks, inbreeding, hybridization, and multiple introductions can all operate within populations of invasive species (Barker et al., 2019; Dlugosch and Parker, 2008; Ellstrand and Schierenbeck, 2000; Hovick and Whitney, 2014; Kolbe et al., 2004). More generally, the impact of demography on levels of genetic variation in invasive populations has been a major focus in the field of invasion genetics since its inception more than 60 years ago (Dlugosch et al., 2015; Lee, 2002); especially with respect to its effect on invasive species’ ability to adapt to novel environments. Under the assumption that invasive species experience strong bottlenecks, and in some cases inbreeding, an erosion of genetic diversity is predicted to reduce fitness and impose constraints on a population’s ability to persist in and adapt to novel environments (Markert et al., 2010; Agashe et al., 2011; Agashe and Bolnick, 2010). The fact that invasive species are frequently able to successfully colonize and adapt to novel environments, despite the negative consequences that a loss of diversity is expected to have on fitness (and by extrapolation, population growth rates), has led to the idea of the “genetic paradox of invasive species” (Allendorf and Lundquist, 2003; Estoup et al., 2016).

While studies have shown that some invasive populations experience a loss of genetic diversity relative to populations in the species’ native range (Grapputo et al., 2005; Michaelides et al., 2018; Tsutsui et al., 2000), these findings are frequently based on a small number of putatively neutral genetic markers. Others have shown that invasive populations can maintain high levels of genetic diversity through processes such as hybridization and multiple introductions stemming from different source populations in the species’ native range (Facon et al., 2008; Lavergne and Molofsky, 2007; Stepien et al., 2005). The paradoxical nature of invasions has therefore been called into question (Dlugosch et al., 2015; Estoup et al., 2016).

At least two general arguments have been made against support for a genetic paradox in invasive species. First, reduced genetic diversity at neutral loci is not equivalent to a reduction in genetic variation that is important for adaptation: invasive populations may show reduced genetic variation at neutral loci but retain variation at loci that are important for adaptation to novel environments (Dlugosch et al., 2015; Estoup et al., 2016). Therefore, populations of invasive species may still show high fitness and an ability to adapt to novel habitats despite a loss of genetic variation at loci that do not underlie fitness-associated traits. Second, invasive populations may not need to adapt to the habitats they are colonizing, therefore removing the paradox altogether (Estoup et al., 2016). For example, populations may adapt to human altered or disturbed environments in their native range, thereby facilitating subsequent range expansions into “anthropogenic” environments (a process termed “anthropogenically induced adaptation to invade”: (Hufbauer et al., 2012)). These arguments describe reasons why successful colonization and adaptation in invasive species may not be paradoxical.

Genetic diversity in neutrally evolving populations of constant size is expected to be proportional to the effective population size and the mutation rate (*θ* = 4*N*_*e*_*μ*; (Watterson, 1975)). Populations of invasive species do not however exist under equilibrium conditions and can experience large fluctuations in population size. Non-equilibrium dynamics such as these are predicted to affect both the amount and type of genetic variation segregating within a population. Genetic drift in small populations can, for example, lead to the fixation of weakly deleterious mutations segregating at low frequencies in source populations (Marsden et al., 2016; Rogers and Slatkin, 2017), and populations that are either increasing (e.g. a growing invasive population) or decreasing (e.g. bottlenecked populations at the front of a range expansion) in size are expected to fix novel beneficial or deleterious mutations, respectively, with higher probability than populations of constant size (Otto and Whitlock, 1997). These theoretical expectations have important implications for the dynamics of adaptation (and maladaptation) during range expansions. We therefore require a better understanding of how changes in population size during biological invasions and range expansion affect genome-wide patterns of diversity and how these changes may alter the average fitness of individuals in invading populations.

When explicit links between genetic variation and adaptive phenotypic variation have not been made, a comparative approach can be used to gain insight into general effects that invasion has on genetic variation. First, genome-wide data can be used to gain a more nuanced understanding of the effects of range expansion on patterns of genetic variation. For example, genetic diversity varies across the genome (Dutoit et al., 2017; Langley et al., 2012) and genomic data from invasive and native populations of an invasive species can be used to test whether genetic diversity is consistently reduced across the genome, or whether certain genomic regions show disproportionately large or small reductions in diversity. Second, while studies of genetic variation in invasive species tend to focus on comparisons between invasive and native populations, comparisons between invasive and closely related non-invasive species can be used to test whether the reduction in genetic diversity observed in an invasive population results in levels of genetic diversity that are below broadly observed levels. Under this approach, non-invasive species are used to quantify the range of naturally-occurring genetic variation and act as a pseudo-control for levels of genetic variation that can be found in viable natural populations.

Quantifying performance across different environments is another way to assess whether range expansions associated with biological invasions have affected fitness in invasive populations. For example, if inbreeding and small population size had led to a decrease in fitness, one prediction is that between-population crosses will display higher fitness than crosses carried out within an inbred population (i.e. heterosis (Oakley et al., 2019)). More generally, if invasive populations suffer from inbreeding depression, we would predict that they will display lower fitness (or some measure correlated with fitness) relative to outbred populations that are found in the native portion of the species’ range (Oakley et al., 2019). Therefore, by combining genome-wide surveys of genetic variation with phenotypic measurements of performance, we can gain a better understanding of the processes affecting genetic variation during biological invasions and how those processes might affect the fitness of individuals in invading populations.

Here, we analyze whole genome sequences collected from 93 individuals sampled across 7 species and 16 populations of African fig fly (genus *Zaprionus*; Figure 1) to test whether invasive populations of *Z. indianus* are outliers with respect to the amount of genetic variation they harbor. We also test whether genetic diversity is affected by genomic features such as the presence of genes and local recombination rates. Finally, we estimate thermal performance curves for 21 populations across 13 species of African fig fly to test whether there is any general relationship between levels of genetic variation and performance across a broad range of temperatures. We find that invasive populations of *Z. indianus* have lower amounts of genetic diversity than populations in their native range, and that both the presence or absence of genes and local recombination rates affect amounts of genetic diversity. However, genetic diversity in invasive populations tends to remain as high or higher than in non-invasive species of *Zaprionus*. Estimates of thermal performance curves indicate that *Z. indianus* has the broadest thermal niche of the species we tested, and despite lower genetic diversity, invasive populations of *Z. indianus* do not show reduced performance relative to populations in their native range. These results suggest that the evolution of a broad thermal niche can facilitate successful range expansion by allowing invasive species to maintain high fitness across a wide range of thermal environments. In turn, large population sizes (and high levels of genetic diversity) in *Z. indianus*’s native range may have acted to buffer invasive populations against critical losses of genetic diversity.

**Figure 1.**
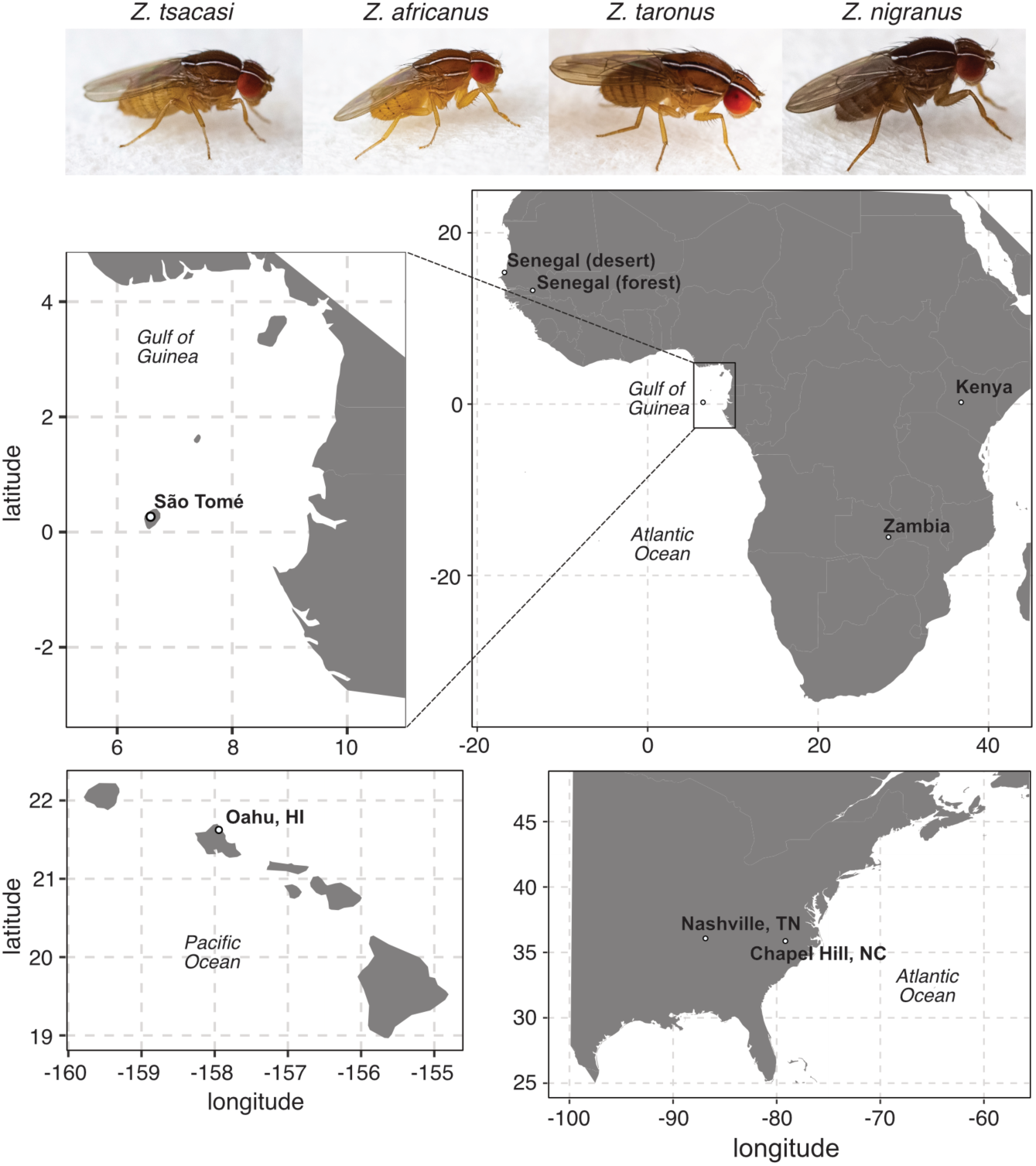
Collection locations across sub-Saharan Africa, eastern North America, and Hawaii. All collection locations are highlighted with bold type. *Zaprionus indianus* was sampled from all locations, *Z. tuberculatus* was sampled from sites in São Tomé and Senegal (forest site), *Z. africanus* from sites in São Tomé and Kenya, and *Z. inermis, Z. tsacasi, Z. taronus*, and *Z. nigranus* from sites in São Tomé. Images of four species of *Zaprionus* are shown for reference.

## Results

### Invasive populations of Z. indianus have less genetic diversity than native populations

A central tenet of the genetic paradox of invasive species is that relatively few individuals colonize invasive parts of their range and that these populations are subject to a loss of genetic diversity, and potentially inbreeding (Allendorf and Lundquist, 2003; Estoup et al., 2016). We tested for relatedness and inbreeding in our samples, but found no evidence that any of the individuals we sampled were closely related (all kinship coefficients estimated within populations < 0.017). There was also no evidence for inbreeding in the invasive range of *Z. indianus* relative to its native range (*F*_1,17_ = 2.37; *P* = 0.14; Figure S1). Below we therefore focus on levels of genetic diversity segregating within populations.

We computed nucleotide diversity across segregating sites (*π*_*SNP*_), the number of segregating sites (*S*), and Tajima’s *D* in non-overlapping 5,000 bp windows across the genome for 16 populations of 7 species of *Zaprionus*. Among populations of the invasive *Z. indianus*, populations sampled in the invasive part of the species’ range display significantly less genetic diversity than populations sampled in their native range: the median number of segregating sites (*S*) in 5 kb genomic windows was between 120 and 160 across North American and Hawaiian populations (5% empirical quantiles: 0 to 10 SNPs; Figure 2; Table S4) and 206 to 233 across African populations (5% empirical quantiles: 21 to 37 SNPs; Figure 2; Table S4). We observed similarly low genetic diversity in invasive, relative to native, populations of *Z. indianus* when we restricted our analysis to genomic windows that overlap an annotated BUSCO gene (Figure S2; Table S5). These results are in line with studies in other systems that have shown lower genetic diversity in invasive relative to native populations of invasive species (Grapputo et al., 2005; Michaelides et al., 2018).

**Figure 2.**
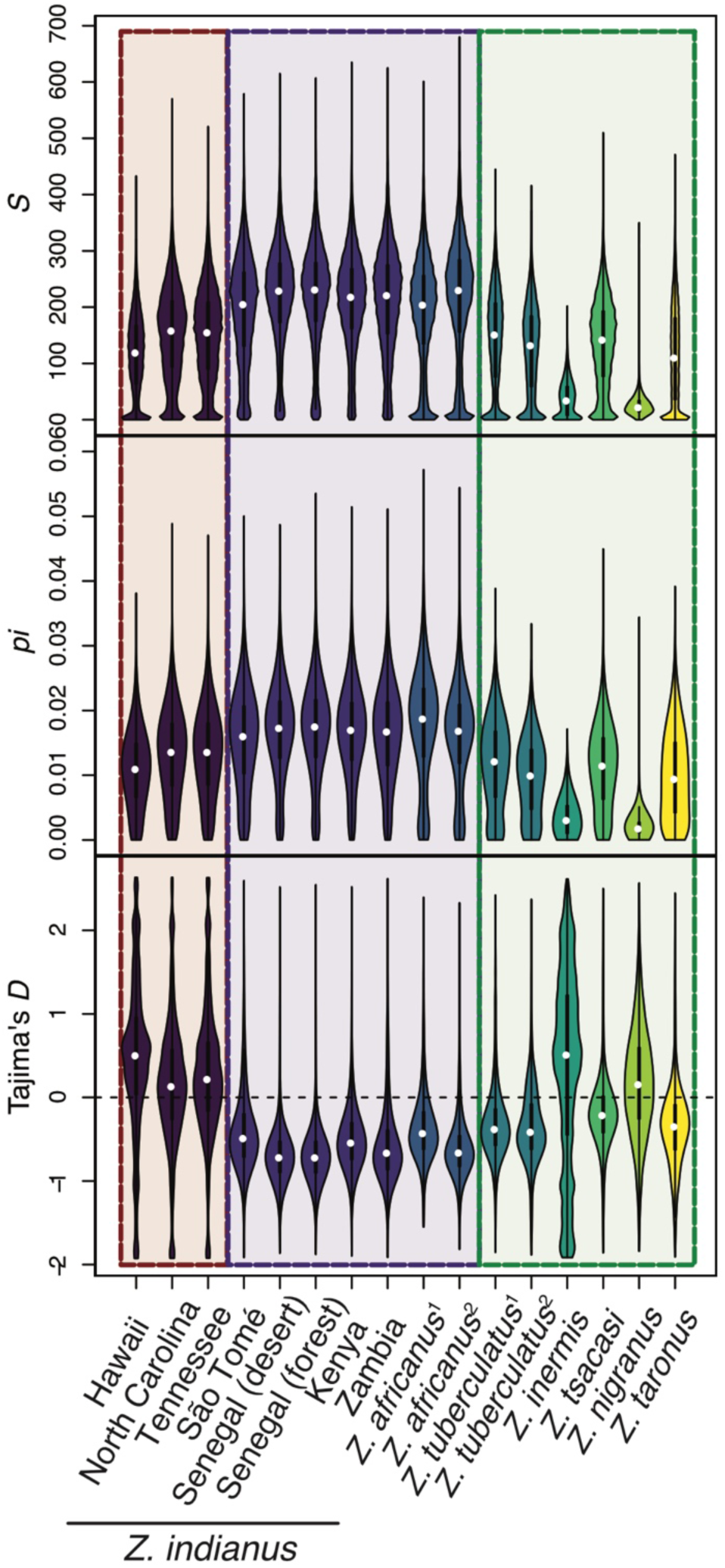
Estimates of genetic diversity summarized across 5 kb genomic windows for each population included in this study. Colored backgrounds group populations as invasive *Z. indianus* (three leftmost violins), native *Z. indianus* and *Z. africanus* (seven central violins), and other species (six rightmost violins). See Figure S2 for estimates for windows overlapping with BUSCO annotations and Table S4 for estimates in all subsamples from populations where we sampled more than four individuals.

Tajima’s *D* also varied across populations of *Z. indianus*, with invasive populations showing positive median genome-wide estimates of Tajima’s *D* (0.13 to 0.5), consistent with a recent contraction in population size, and native populations showing negative median estimates of Tajima’s *D* (−0.48 to −0.72), consistent with range expansion (Figure 2; Table S4).

### *Genetic diversity in invasive populations of* Z. indianus *is not exceptionally low*

While levels of genetic diversity were reduced in invasive populations of *Z. indianus*, we found that these populations still harbor as much, or more, genetic diversity than naturally occurring non-invasive species of *Zaprionus* (Figure 2). For example, *S* within populations of *Z. tuberculatus* and *Z. tsacasi* (median *S* = 141 and 143, respectively) is comparable to *S* in invasive populations of *Z. indianus* (median *S* = 120 to 140). By contrast, populations of *Z. nigranus, Z. taronus*, and *Z. inermis* all show markedly less genetic diversity (median *S* = 23, 106, and 35, respectively) than any of the *Z. indianus* populations we sampled. The broadly distributed species *Z. africanus* (the closest relative to *Z. indianus* included in this study) showed amounts of genetic diversity comparable to those in native populations of *Z. indianus*. Similar to comparisons among populations of *Z. indianus*, we recovered the same relationship when restricting our analysis to genomic windows that overlap an annotated BUSCO gene (Tables S4 and S5). Patterns of genetic diversity across populations of *Z. indianus* and other species of *Zaprionus* suggest that, under the assumption that the amount of genetic diversity in non-invasive species is a reasonable proxy for the amount of genetic diversity a population can have without negative fitness effects, high levels of genetic diversity in the native range of invasive species may buffer invasive populations against losses of diversity below levels that would negatively affect fitness or adaptive potential.

### Genetic diversity is affected by genome architecture

While population bottlenecks occurring in invasive populations are expected to lower genetic diversity broadly across the genome (Ellegren and Galtier, 2016; Hyten et al., 2006), other processes can act locally within the genome and either reduce (e.g. selective sweeps; (Charlesworth et al., 1997)) or maintain genetic diversity (e.g. balancing selection; (Aguilar et al., 2004; Charlesworth, 2006; Lindtke et al., 2017)). The fitness consequences of diversity across loci is also not expected to be equal. Genetic diversity therefore varies greatly across the genome and the interaction between selection and recombination rate can lead to a genome-wide “landscape” of genetic diversity that is correlated with aspects of genome architecture such as recombination rates or gene density (Begun and Aquadro, 1992; Burri et al., 2015; Dutoit et al., 2017; Nachman, 2001). We generated gene annotations and used estimates of recombination rates to test the relationship between genetic diversity in invasive and native populations of *Z. indianus* and these features of the genome.

Across all genomic windows, we found that the amount of genetic diversity (*S*) within a given genomic window varied depending on whether that window overlapped-with, was adjacent-to, or was distant-from an annotated gene (population level comparisons: GLMs: all *P* < 0.0001). Genetic diversity tended to be lower in windows that overlap an annotated gene compared to windows that were either within or outside of 5kb from an annotated gene (12 of 15 populations; binomial test: *P* = 0.035; Figure S3). Three populations did not follow this trend: the two *Z. africanus* populations and the *Z. tuberculatus* population sampled from Senegal. For populations that had less diversity in windows that overlapped an annotated gene, mean diversity tended to be 3.5 to 21.5% lower than in windows within 5kb of an annotated gene, and 0.6 to 32.9% lower than in windows further than 5kb from an annotated gene (Table S6). We did not, however, observe an interaction between invasion status (invasive *Z. indianus*, native *Z. indianus*, or non-invasive *Zaprionus*) and window-location relative to annotated genes and the median amount of genetic diversity (*S*) observed across windows (GLM: *P* = 0.99). The amount of genetic diversity within a genomic region is therefore affected by the presence (or absence) of genes, but broad scale differences in diversity between “genic” and “non-genic” regions is not systematically altered -for example, due to selection preferentially maintaining genetic diversity in or around genes -during the course of invasion.

We next explored the relationship between genetic diversity (*S*) and mean estimates of population-scaled recombination rates in 5kb genomic windows (Materials and Methods). Consistent with previous studies (Begun et al., 2007; Begun and Aquadro, 1992; Burri et al., 2015; Dutoit et al., 2017; Samuk et al., 2017), *S* was positively correlated with recombination rate across genomic windows in all *Z. indianus* populations (all Spearman’s *ρ* > 0.5; Figure 3a). The strength of this correlation did not systematically differ between invasive and native populations of *Z. indianus* (linear model: *F*_1,12_ = 0.1025; *P* = 0.7543); however, the mean difference in *S* between invasive and native populations of *Z. indianus* was correlated with local recombination rate (analysis of windows with mean *S* between 150 and 300 in *Z. indianus*’s native range; Figure 3b). This correlation was relatively weak when considering the raw difference in *S* (Spearman’s *ρ* = 0.071; Figure S4), but was stronger when the difference in *S* was scaled by mean *S* for genomic windows binned by recombination rate quantiles (Spearman’s *ρ* = 0.231; Figure 3b). This finding shows that during invasions, the loss of diversity is not uniform across the genome, but is greater in genomic regions experiencing lower recombination rates.

**Figure 3.**
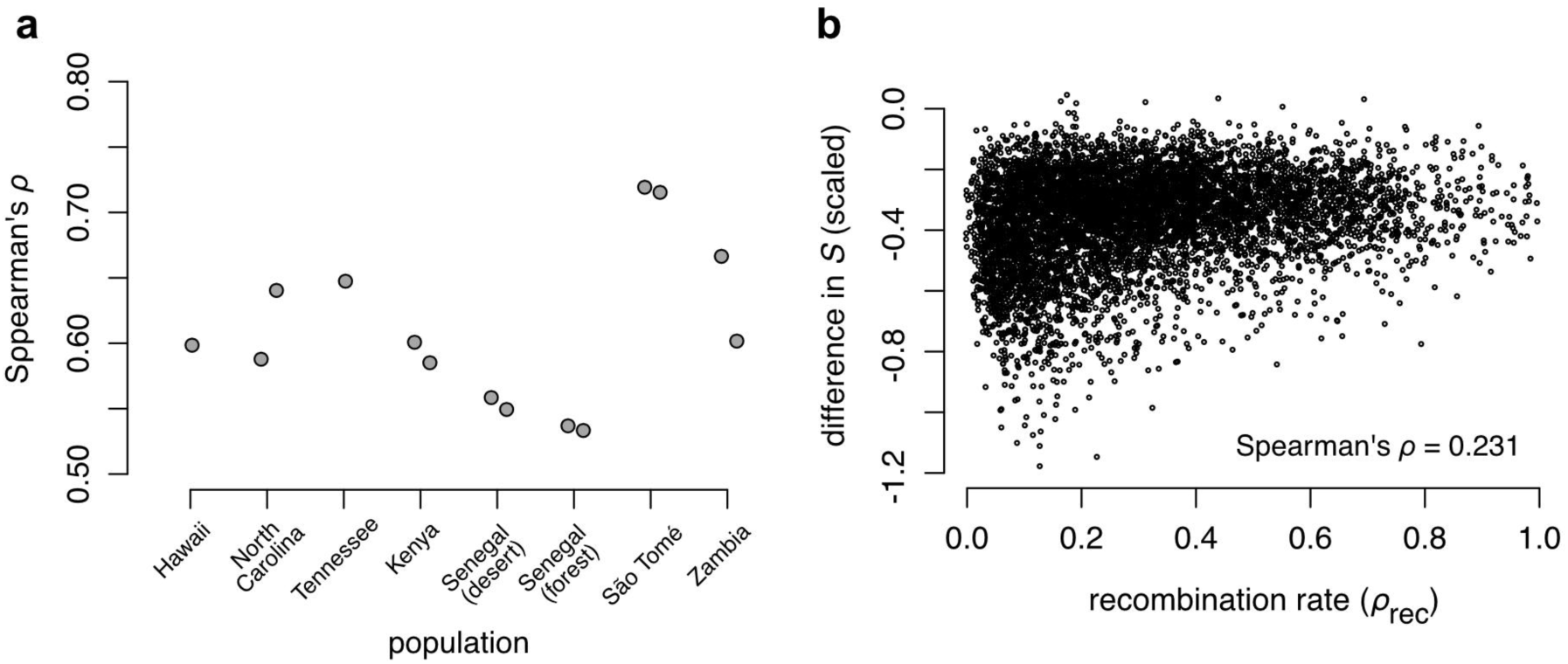
Correlations between genetic diversity (the number of segregating sites: *S*) and recombination rate across populations of *Z. indianus* (**a**). Correlations did not systematically differ between populations in the invasive and native regions of the species’ range. However, the mean difference in diversity for a given genomic window was positively correlated with recombination rate (**b**). Populations with two points in panel **a** represent populations where we sampled more than 4 individuals and estimated *S* using two independent random subsamples of those individuals. In panel **b**, the difference in *S* (mean in invasive populations -mean in native populations) was scaled by mean levels of diversity for a given recombination rate quantile.

### Invasion alters the genome-wide “landscape” of genetic diversity

As a second test of how genetic diversity is distributed across the genome and whether that genome-wide distribution is altered during biological invasions, we compared genetic diversity between species in genomic windows that spanned annotated BUSCO genes. Consistent with a genome-wide landscape of genetic diversity, we found that genetic diversity (*S*) within annotated BUSCO genes was correlated between populations and species of *Zaprionus* (Figure 4; see Figure S5 for randomization test). As expected, correlations in *S* were weaker for interspecific comparisons than for intraspecific comparisons (Figure 4a). The strongest interspecific correlation we observed was between *Z. indianus* from Zambia and *Z. africanus* from São Tomé (Spearman’s *ρ* = 0.4504; Figure 4b) and the weakest was between *Z. inermis* from São Tomé and *Z. africanus* from Kenya (Spearman’s *ρ* = 0.1888; Figure 4c). Across all pairwise comparisons, there was a significant negative relationship between the correlation in *S* across the genome and the genetic distance between the species being tested (Mantel *r* = −0.816; *P* = 0.0001; Figures 5a and b). This trend was even stronger among comparisons between more closely related species pairs (i.e. “within-clade” comparisons; Mantel *r* = −0.83; Figure 5c). The pattern of weaker correlations in *S* between more genetically diverged species is consistent with evolution of aspects of genome architecture (e.g. intron size, recommendation rates, and / or transposable element evolution) altering the genome-wide landscape of diversity; however, explicit tests of these mechanisms are still needed.

**Figure 4.**
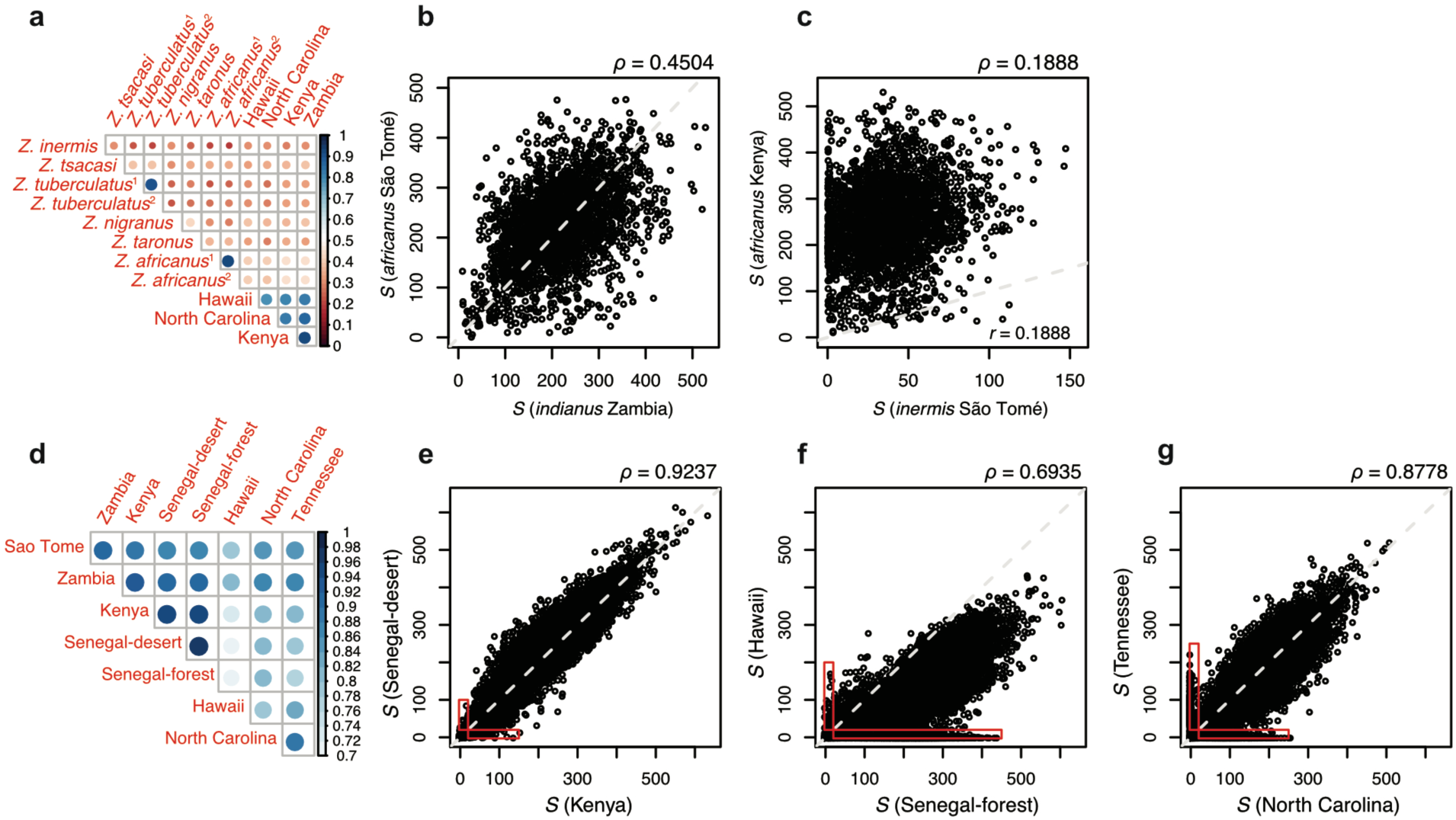
Genetic diversity (*S*) is correlated across regions of the genome containing annotated single-copy orthologs in interspecific comparisons (**a**). The closely related species *Z. africanus* from São Tomé and *Z. indianus* from Zambia had the strongest interspecific correlation in *S* (**b**; see Figure 5a for phylogeny) and the distantly related species *Z. africanus* from Kenya and *Z. inermis* from São Tomé had the weakest correlation in *S* (**c**). Genetic diversity is strongly correlated in all pairwise comparisons between populations of *Z. indianus* (**d**). Panels **e** and **f** show data from the strongest and weakest between-population correlations for *Z. indianus*, and panel **g** shows the strongest between invasive populations of *Z. indianus*. Red rectangles in panels **e** through **g** highlight genomic windows that have low diversity in one population (fewer than 10 segregating SNPs), but higher diversity in the other (more than 10 segregating SNPs).

**Figure 5.**
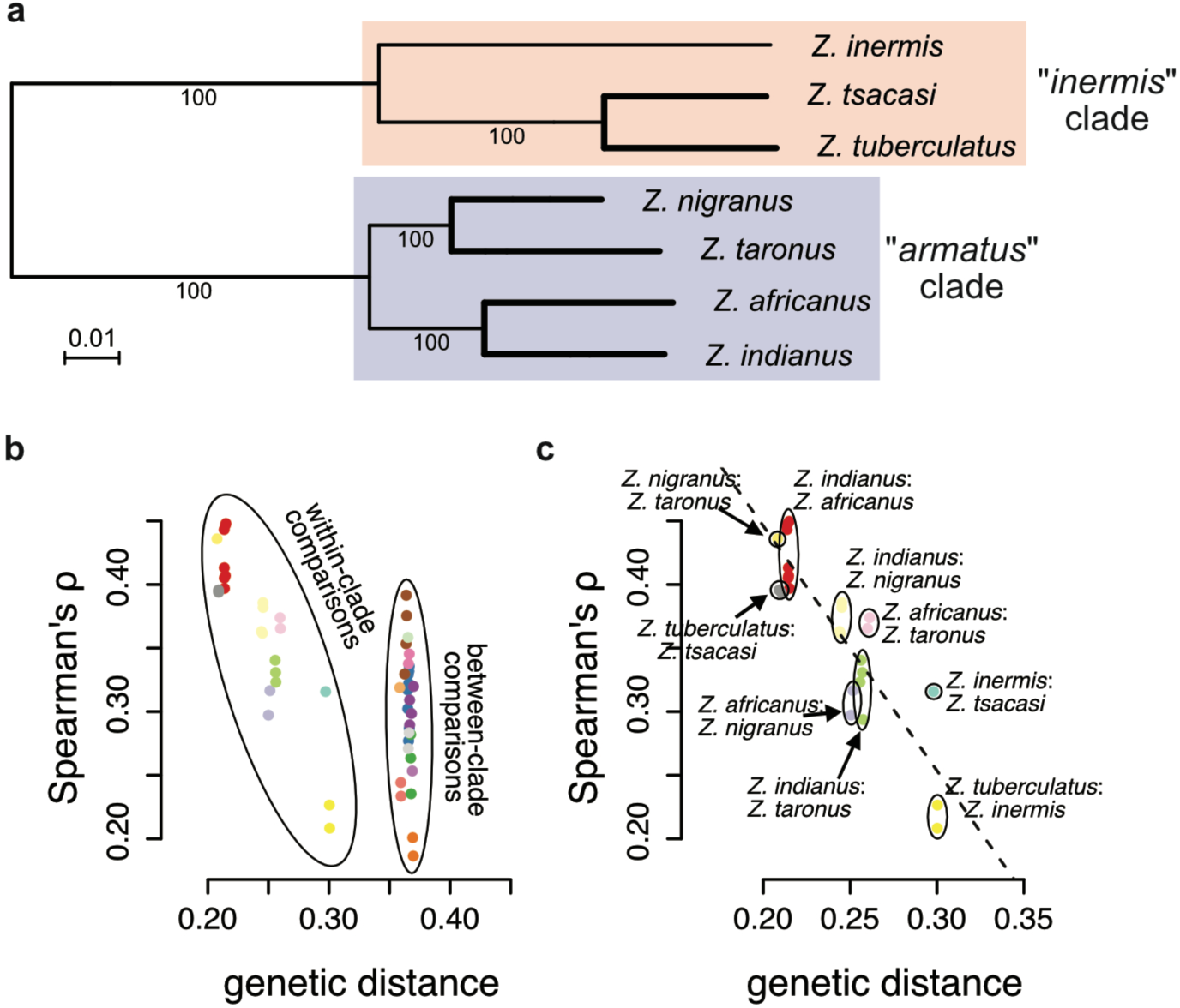
The correlation of genetic diversity (*S*) across the genomes of different species (**a**) decreases with increasing genetic distance (**b** & **c**). Maximum likelihood phylogeny in **a** was estimated with RAxML run on an alignment of 1709 BUSCO genes annotated across all seven species genomes. Correlation coefficients decreased with increasing genetic distance (Mantel *r* = −0.816; **b**), with this pattern being particularly evident for within-clade comparisons (highlighted in **c**).

As with interspecific comparisons restricted to BUSCO windows, genetic diversity (*S*) was significantly correlated across all genomic windows between populations of *Z. indianus* (Figure 4d). Among African populations of *Z. indianus*, this was not a function of how closely related two populations were: geographically distant populations in Senegal and Kenya or Senegal and Zambia (Figure 1) show among the strongest correlations in *S* (Spearman’s *ρ* > 0.885; Figures 4d & 4e). In general, the correlation in *S* was lower when a native and an invasive population were being compared versus comparisons made among native populations (maximum *ρ* = 0.865, minimum *ρ* = 0.694; Figure 4d & 4f). Among invasive populations, North Carolina and Tennessee populations showed the strongest correlation in *S* (Spearman’s *ρ* > 0.8778; Figure 4g); however, this correlation was lower than the five strongest correlations, all observed between African populations.

In contrast to *S*, Tajima’s *D* was either weakly correlated or uncorrelated between species (Figure 6a), and there was no relationship between the strength of correlation in Tajima’s *D* and phylogenetic distance (*P* > 0.1). By contrast, Tajima’s *D* was significantly correlated across genomic windows between all populations of *Z. indianus* (*P* < 0.001; Figure 6b). However, Tajima’s *D* was more weakly correlated than *S* (compare Figure 6b to Figure 4d). As with *S*, the strongest correlations in Tajima’s *D* were between populations sampled in Africa, consistent with a shared demographic history in the native part of *Z. indianus*’ range (Figure 6b and 6c). By contrast, Tajima’s *D* was only weakly correlated in between-continent comparisons (strongest between-continent *ρ* = 0.192; weakest *ρ* = 0.0167; Figure 6d) and moderately correlated between the two populations sampled in the eastern USA (Tennessee and North Carolina; *ρ* = 0.2002; Figure 6e). The fact that Tennessee and North Carolina have both been recently colonized during *Z. indianus*’s expansion into North America (∽20 years ago; (Gibert et al., 2016)), yet show relatively modest correlations in Tajima’s *D*, illustrates how demographic events can rapidly alter the frequency of alleles across the genome and suggests that these two locations are experiencing semi-independent colonizations, rather than an expansion driven by a single panmictic population.

**Figure 6.**
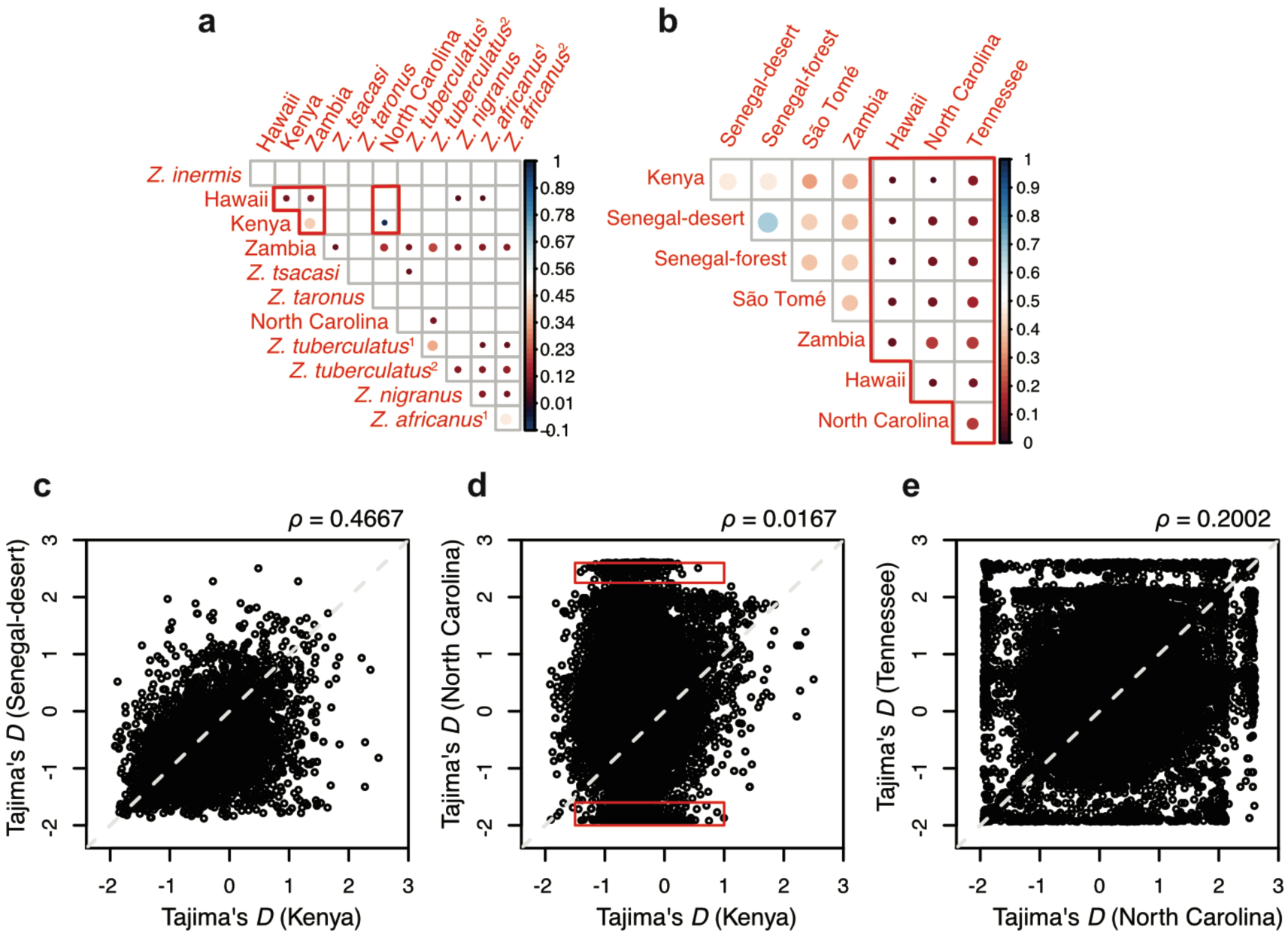
Tajima’s *D* is weakly correlated or uncorrelated between different species of *Zaprionus* (**a**) and populations of *Z. indianus* (**b**). The strongest correlations in S (outside of populations of *Z. indianus* sampled from two locations in Senegal) were between geographically distant populations of *Z. indianus* in its native Africa (**c**) and the weakest correlation between *Z. indianus* populations was between a native and an invasive population (**d**). The two geographically proximate invasive populations of *Z. indianus* in North America showed a moderate correlation in Tajima’s **D** across windows. Red polygons in **a** highlight comparisons between populations of *Z. indianus* and in **b** highlight comparisons between invasive populations. Red rectangles in panel **d** highlight genomic windows that show a pronounced shift in Tajima’s *D* between the invasive population in North Carolina and the native population in Kenya.

The correlation of genetic diversity (*S*) between *Z. indianus* and other species of *Zaprionus* (Figures 4 and 5) allowed us to test whether this correlation is maintained following biological invasion. We find that interspecific correlations in *S* are weaker between invasive populations and other species than between native populations and other species (6 of 8 species/population comparisons; Fisher’s exact test: *P* = 0.0093; Figure 7). When we restrict this same analysis to include one population per species, the correlation in genetic diversity remains weaker between invasive populations of *Z. indianus* and other *Zaprionus* species than between native populations of *Z. indianus* and other *Zaprionus* in four of the 6 species-level comparisons (Fisher’s exact test: *P* = 0.03). Native populations of *Z. indianus* also never showed consistently weaker correlations than invasive populations (Figure 7). These findings illustrate how demographic and selective processes associated with biological invasions do not only result in a general reduction in diversity in invasive populations, but also alter the genome-wide distribution of that diversity.

**Figure 7.**
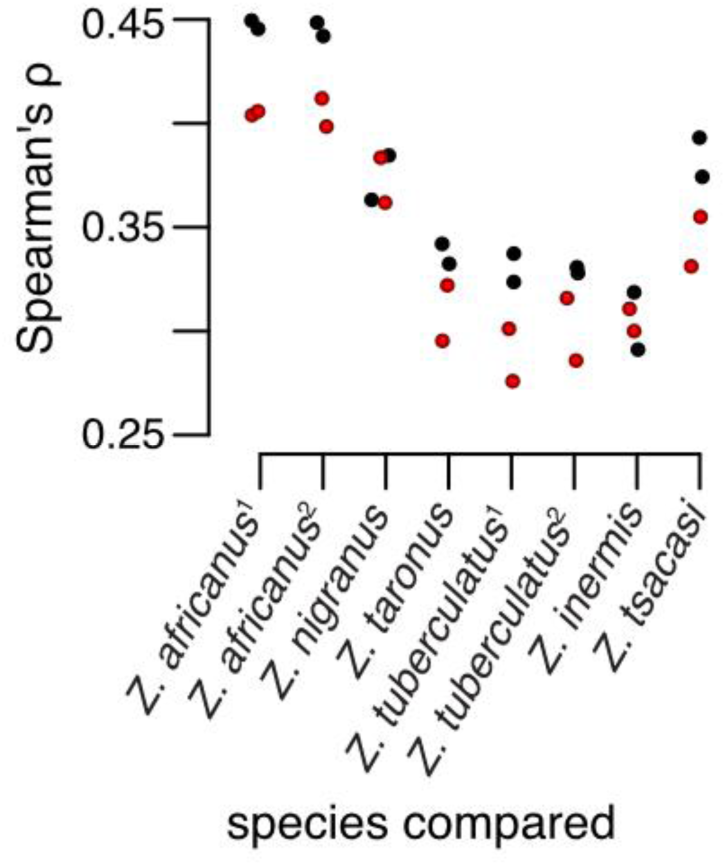
The correlation in genetic diversity (*S*) between invasive populations of *Z. indianus* and other species of *Zaprionus* (dark red points) is weaker than the correlation in *S* between native populations of *Z. indianus* and other species of *Zaprionus* (black points).

### Genetic diversity is correlated with thermal niche breadth

Genetic diversity can, in some cases, be positively correlated with measures of performance or fitness (Briskie and Mackintosh, 2004; Leimu et al., 2006; Markert et al., 2010; Reed and Frankham, 2003). We therefore tested for relationships between genetic diversity and fitness/performance, as measured from thermal performance curves.

We estimated thermal performance curves using a hierarchical Bayesian framework ((Tittes et al., 2019); SI). Parameters summarizing performance were estimated at the population level, allowing us to make comparisons between populations of *Z. indianus* sampled from three populations from their invasive range in eastern North America (Florida, North Carolina, and New York, USA) and their native range in Africa (São Tomé, Senegal, Kenya, and Zambia).

Among populations of *Zaprionus* species with estimates of genetic diversity and thermal performance, thermal niche breadth (*B*_*50*_) and the total area of the thermal performance curve (*A*_*c*_) were positively associated with genetic diversity (Figure 8a; linear models: *B*_*50*_: *F*_1,11_ = 10.1; *P* = 0.009; *A*_*c*_: *F*_1,11_ = 7.402; *P* = 0.020; note that *A*_*c*_ and maximum fitness parameters were strongly correlated; *r* = 0.99). By contrast, there was no relationship between levels of genetic diversity and thermal optimum (*F*_1,11_ = 1.625, *P* = 0.23). Linear models fit to our complete data set should however be interpreted with caution because of phylogenetic nonindependence and uneven sampling across species (e.g. we sampled multiple populations of *Z. indianus, Z. africanus*, and *Z. tuberculatus*). We therefore also fit models that only included a single population from each of the seven species sampled from São Tomé (Figure 8b). Using this reduced data set, we still find support for a positive relationship between genetic diversity and thermal niche breadth (*B*_*50*_: *F*_*1,5*_ = 8.244; *P* = 0.035; *R*^2^ = 0.55), but no relationship between genetic diversity and *A*_*c*_ or thermal optimum (both *P* > 0.1). These results are consistent with genetic diversity being higher in species that are able to exploit a broad range of thermal environments, potentially due to these species maintaining large population sizes.

**Figure 8.**
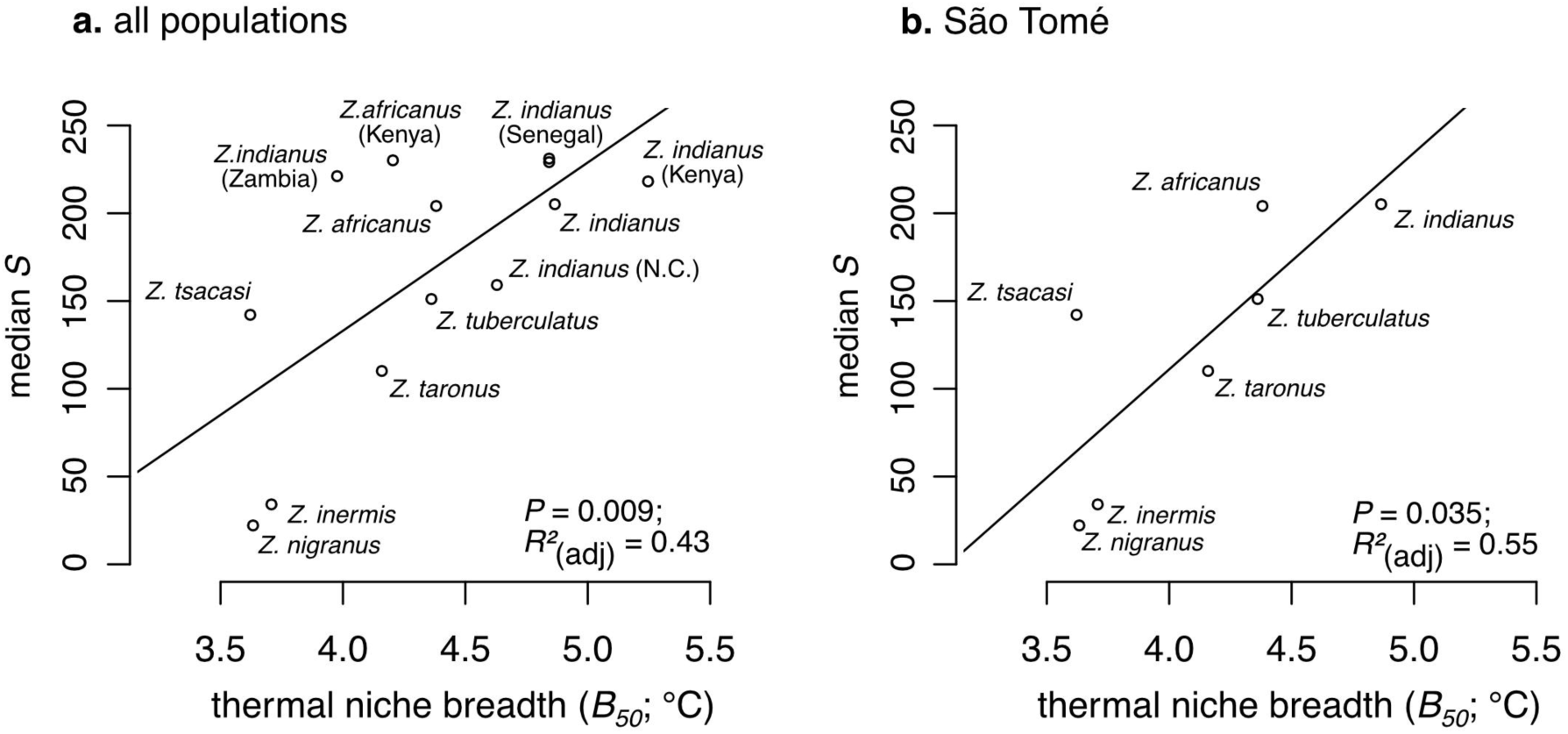
Populations and species with a larger thermal niche breadth (*B*_*50*_) also harbor more genetic diversity (median number of segregating sites; *S*). Results for all populations (**a**) and for only the seven species sampled on the island of São Tomé (**b**) are shown. Species names that are not labeled with a location in panel (**a**) are from São Tomé. Results from linear models testing the relationship between thermal niche breadth and genetic diversity are reported in the bottom right of each panel.

### Performance is not reduced in invasive populations

Among the seven species we collected population genomic data from, *Z. indianus* tended to have the highest estimated *B*_*50*_ (Figure 8). We tested the generality of this pattern by estimating thermal performance curves for an additional 6 species and 13 populations of *Zaprionus*. In this data set, populations of *Z. indianus* from both their native and invasive ranges consistently have larger estimates of *B*_*50*_ than other populations and species (Figure 9). Two exceptions to this trend are that the *B*_*50*_ of *Z. indianus* from Zambia is more similar to *Z. africanus* than other populations of *Z. indianus*, and the *B*_*50*_ of *Z. gabonicus* is similar to populations of *Z. indianus* (Figure 9).

**Figure 9.**
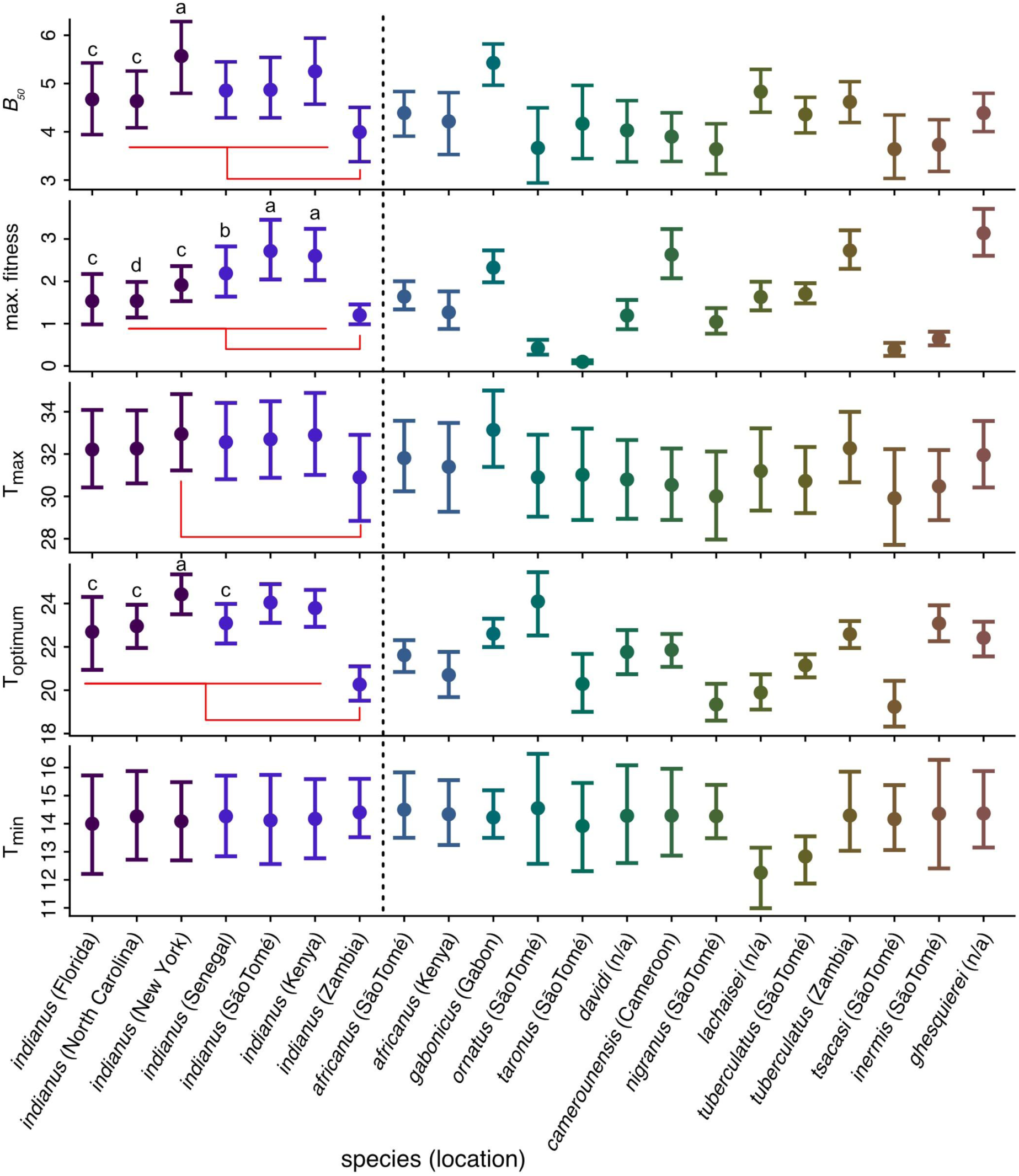
Median and 95% credible intervals for parameters estimated by jointly fitting performance curves to thermal performance data collected at mean temperatures of 13.9 to 28.9°C. Populations of *Z. indianus* are shown to the left of the dashed line and all other species of *Zaprionus* to the right. For populations of *Z. indianus*, parameter estimates that had a probability greater than 0.95 of being different from one another in pairwise comparisons (analogous to *P* < 0.05) are denoted by letters (‘a’ > ‘c’ and ‘d’, and ‘b’ > ‘d’). Red bars indicate significant differences between the population of *Z. indianus* from Zambia and other populations of *Z. indianus*.

We next tested whether invasive and native populations of *Z. indianus* differed in maximum fitness, as measured by the total number of adult progeny produced by a single pair of flies, at their optimal temperature (“stretch” parameter reported in Tittes et al. 2019; “max. fitness” in Figure 9). African populations of *Z. indianus* from Kenya and São Tomé have higher maximum fitness than invasive populations from Florida, New York, and North Carolina, while the population from Senegal only had higher maximum fitness than the invasive population sampled from North Carolina (Figure 9). All populations of *Z. indianus* other than those from Florida and North Carolina had higher maximum fitness than the population from Zambia (Figure 9). The lower maximum fitness observed in invasive populations, compared to populations from Kenya and São Tomé, could be explained by a loss of genetic diversity having negative effects on fertility, fecundity, or survivorship. However, if a loss of genetic diversity underlies a decrease in maximum fitness, one prediction is that these effects would be particularly pronounced at stressful temperatures (i.e. near the minimum and maximum acceptable temperatures), thereby reducing the breadth of the performance curve. This was not the case, as estimates of thermal minima and maxima did not consistently differ between invasive and native populations (T_min_ and T_max_ parameters, respectively; Figure 9). There was also no evidence that maximum fitness in invasive populations (Figure 9) was different from that in the African populations collected in Senegal, despite the latter having high levels of genetic diversity. Therefore, we did not find consistent evidence that the loss of diversity associated with *Z. indianus*’s range expansion into North America has reduced the generally high performance that this species displays across a broad range of temperatures.

In addition to comparisons among populations of *Z. indianus*, populations of *Z. indianus* from Kenya and São Tomé both had maximum fitness equal-to or greater than the 12 other species of *Zaprionus* we measured (Figure 9). The population from Senegal had a similarly high estimate of maximal fitness compared to all other species, except *Z. ghesquierei*, which showed a higher maximal fitness than all three invasive North American populations of *Z. indianus*, as well as *Z. indianus* from Senegal and Zambia (Figure 9). Three other species were estimated to have a maximal fitness that was greater than *Z. indianus* from at least two of the three North American populations: *Z. gabonicus, Z. camerounensis*, and *Z. tuberculatus*. This result suggests that the range of temperatures over which a species or population can maintain relatively high fitness (e.g. B_50_) is a more important factor differentiating *Z. indianus* from non-invasive species of *Zaprionus* than the absolute maximal fitness/performance that individuals in a population can achieve.

## Discussion

Identifying the genetic changes that are associated with biological invasions, along with effects those changes have on performance and fitness, is important for understanding the evolutionary controls acting on invasive species (Estoup et al., 2016; Lawson Handley et al., 2011; Lee, 2002; Stapley et al., 2015). We have shown that the range expansion of the invasive African fig fly, *Z. indianus*, has resulted in a loss of diversity in invasive populations, but that levels of diversity in invasive populations remain as high or higher than those observed in other species in the genus *Zaprionus*. We also show that invasive and native populations of *Z. indianus* do not differ in their ability to maintain high performance across a broad range of temperatures. This finding is in contrast with studies that have found a positive correlation between genetic diversity and measures of fitness ((Leimu et al., 2006; Markert et al., 2010; Reed and Frankham, 2003); but see (Lammi et al., 1999)).

A number of evolutionary mechanisms that act to increase (or maintain) genetic diversity in invasive populations could help explain the moderate levels of genetic diversity in invasive populations of *Z. indianus*. For example, multiple introductions from distinct populations in the native range and admixture can both act to generate high levels of genetic diversity in invasive populations (Stepien et al., 2005; Dlugosch and Hays, 2008; Rosenthal et al., 2008; Barker et al., 2017). The fact that invasive populations of *Z. indianus* harbour as much or more genetic diversity than populations of other *Zaprionus* species suggests that processes such as multiple introductions, a large number of founding individuals, or admixture may be acting to maintain moderate levels of genetic diversity in these populations. High levels of genetic diversity in native populations of *Z. indianus* (Figure 2) may also act to buffer invasive populations against a loss of diversity, and may help explain the moderately high level of diversity remaining in invasive populations of *Z. indianus*.

The broad thermal niche of *Z. indianus* populations across their native and invasive ranges may also help to explain this species’ high level of genetic diversity and success as an invasive species. For example, if *Z. indianus* evolved a broad thermal niche prior to expanding their range out of Africa, this could have led to large and broadly distributed populations that would build up high levels of genetic diversity. If alleles underlying thermal performance traits became fixed (or present at a high frequency) in native populations prior to range expansion, reductions in genetic diversity would not remove those high-frequency adaptive alleles. This scenario is related to the idea that adaptation to anthropogenically disturbed environments in the native portion of a species range can facilitate subsequent range expansions (Lee and Gelembiuk, 2008; Hufbauer et al., 2012). While the timing of adaptive trait evolution in populations of *Z. indianus*, relative to their range expansion, is not know, anecdotal evidence from collecting *Zaprionus* across locations in their native Africa suggest that *Z. indianus* possess a generalist and ‘invasive-like’ life history (Pers. Obs.). For example, we have collected *Z. indianus* near cities or rural human settlements in rainforest, savannah, and desert environments, and *Z. indianus* is among the first and most abundant species to visit fruit traps across these environments. If the traits that underlie a broad thermal niche evolved in African populations prior to range expansion, then lower diversity in invasive populations may not have negatively affected the fitness of *Z. indianus* in the invasive part of their range.

Estimates of thermal performance curves suggest that, despite the climate being markedly cooler at sample sites in northeastern North America (e.g. North Carolina and New York) than at African sites, populations of *Z. indianus* in North America have not evolved to tolerate more temperate climates that African populations (estimated minimum temperature where fitness falls to 0 does not differ between any population of *Z. indianus*; T_min_ parameter in Figure 9). Thermal niche breadth also does not systematically differ between *Z. indianus* from New York, São Tomé and Kenya, and these populations have significantly wider thermal niches than almost all other *Zaprionus* species or population in our data set (all *P* < 0.05 in pairwise comparisons; Figure 9). Together, these results indicate that the thermal niche of North American and African *Z. indianus* has not evolved to be different, and that the loss of genetic diversity in invasive populations has not reduced the broad range of temperatures across which *Z. indianus* is capable of maintaining relatively high fitness (i.e. *B*_*50*_; Figure 9).

An alternate interpretation for the lack of differences in thermal performance curves between invasive and native populations of *Z. indianus* is that the lower level of genetic diversity in North American populations of *Z. indianus* is constraining potentially adaptive niche evolution in these populations. The use of other *Zaprionus* species as “pseudo”-controls for levels of diversity that segregate in natural populations suggest this is unlikely; invasive populations of *Z. indianus* still harbour amounts of genetic diversity similar to or larger than populations of other *Zaprionus* species. However, future work testing levels of additive genetic variation for traits involved in thermal tolerance are needed to quantify the adaptive potential of the different populations and species.

We were also able to use different species of *Zaprionus* to show that there is a generally conserved genomic landscape of diversity that decreases with increasing phylogenetic distance (Figures 5b and 5c). Correlated levels of genetic diversity have been shown between a pair of divergent bird species (Dutoit et al., 2017), but to the best of our knowledge, this has not been tested in a phylogenetic context until now. We used the correlation in diversity between species to show that the genome-wide landscape of genetic diversity is altered in invasive, relative to native, populations of *Z. indianus* (Figure 7). We also showed that the difference in the amount of genetic diversity between invasive and native populations of *Z. indianus* tends to be larger in regions of the genome with low recombination rates (Figure 3b). Previous studies have reported a similar pattern with respect to lower genetic diversity on the X-chromosome in non-African populations of the fruit fly *Drosophila simulans* (Begun and Whitley, 2000; Schöfl and Schlötterer, 2006). We have yet to identify the scaffolds that compose the sex chromosomes in the genome assemblies we report here, but larger reductions in diversity on the X-chromosome relative to the autosomes is likely to contribute to the pattern of reduced genetic diversity in regions of low recombination. One mechanism that could explain this pattern is linked selection (Begun and Whitley, 2000; Charlesworth et al., 1997, 1997; Nordborg et al., 1996), as selection will have a larger spatial effect on levels of genetic diversity in regions of the genome with low recombination rates. Future work is needed to identify both the chromosome structure of *Zaprionus* species and regions of the genome potentially subject to selection in invasive populations of *Z. indianus*.

If regions of the genome containing alleles that control phenotypes associated with adaptation to climate lose diversity in invasive populations (either due to selection or demographic events), those populations may become constrained with respect to their ability to adapt to novel climatic conditions. As mentioned above, we ultimately need to quantify levels of additive genetic variation in thermal performance related traits to better understand the relationship between genetic diversity and a population’s ability to adapt to novel climates (Chevin, 2013; Dlugosch and Parker, 2008; Franks and Hoffmann, 2012; Hoffmann and Sgrò, 2011).

Traits associated with thermal performance are only one suite of traits potentially under selection during colonization and range expansion into novel environments. Other traits such as competitive ability (Blossey and Notzold, 1995) and the ability to utilize a broad range of other habitats (Lee and Gelembiuk, 2008) can affect the success of invasive species. Of particular note is the fact that we quantified thermal performance on a single food resource (standard cornmeal media for *Drosophila*). Measuring performance in the same populations and species analyzed here, across a wide range of diets, would be useful to determine the extent to which invasive populations of *Z. indianus* have adapted to successfully colonize the broad range of environments they now occupy.

In conclusion, we have shown that genetic diversity in invasive populations of *Z. indianus* is lower than in native populations (Figure 2), and that the genome-wide distribution of genetic variation is perturbed in invasive populations (Figures 3b, 4, 5, and 7). These results illustrate how range expansions associated with invasion can alter levels of genetic variation across the genome. Despite these effects of invasion on genetic diversity, invasive populations of *Z. indianus* maintain as much or more genetic diversity than non-invasive congeneric species (Figure 2), and both invasive and native populations of *Z. indianus* are capable of maintaining high fitness across a broad range of thermal environments (Figures 8 and 9). These results show how measures of fitness in invasive species can be decoupled from amounts of genetic diversity. They also suggest that adaptation to an “invasive” niche prior to range expansion may be a more important event in the evolutionary history of an invasive lineage than the demographic events that take place during subsequent expansion of their range into novel environments. Future work identifying the evolutionary history of invasive traits (and their underlying genetic variation) will be central to our understanding of the factors controlling the success and spread of invasive species.

## Material and Methods

### Population sampling

We sampled wild populations of Drosophilids at five locations across the native range of *Z. indianus* in sub-Saharan Africa and three locations in the invasive range in North America and Hawaii (Figure 1) using traps baited with bananas (SI). To assess patterns of genetic diversity across populations of *Z. indianus* and non-invasive species of *Zaprionus*, we focus on *Z. indianus* populations from Kenya, Zambia, São Tomé, and Senegal (both forest/savannah outside of Niokolo-Koba National Park and coastal desert in the North West) in their native Africa, and Hawaii (Oahu), Tennessee (Nashville), and North Carolina (Chapel Hill) in their invasive range. Across African locations we also sampled two populations of *Z. africanus* (São Tomé and Kenya), two populations of *Z. tuberculatus* (São Tomé and the Senegal-desert site), and one population each of *Z. inermis, Z. tsacasi, Z. taronus*, and *Z. nigranus* (São Tomé).

### Genome assembly and annotation

To facilitate population genomic analyses, we generated genome assemblies *de novo* for each of the seven *Zaprionus* species described above. Genomes were assembled using data generated from Illumina and Nanopore sequencers (SI). We used the BUSCO annotation pipeline (Waterhouse et al., 2018) to assess assembly quality and quantify the presence of—and generate annotations for—2,799 Benchmarking Universal Single-Copy Orthologs that have been curated in 25 different species of Diptera. The percentage of single-copy BUSCOs annotated in a genome assembly provides an estimate of how well that assembly covers gene space while the number of duplicated BUSCOs is a useful metric for identifying the prevalence of misassemblies, for example, due to heterozygous alleles being assembled into distinct contigs (Pryszcz and Gabaldón, 2016; Waterhouse et al., 2018). Finally, we generated gene annotations using RNA sequence data and the `MAKER` annotation pipeline (Campbell et al., 2014) for 6 of the 7 *de novo* genome assemblies (we did not collect RNA-seq data for *Z. inermis*; SI). Our sequencing, assembly, and annotation approaches resulted in genome assemblies with scaffold N50s between 336 kb and 2.45 Mbp, complete single-copy BUSCO annotations of 90.7 to 97.4%, and 9,275 to 11,071 annotated transcripts (Table S2).

### Population resequencing and genotyping

To quantify levels of diversity segregating within populations of *Zaprionus*, we generated resequence data from 93 individuals across 16 populations and 7 species (minimum N = 3; maximum N = 11; Table S3). We extracted genomic DNA from either individual wild-caught flies or from a single offspring of a wild-caught female (i.e. first generation offspring) using Genetra Puregene Tissue Kits (Qiagen, Valencia, CA, USA), constructed barcoded libraries for sequencing using KAPA HyperPrep kits (Roche Sequencing, Pleasanton, CA) with a target fragment size of 500 bp, and sequenced in pools of 10 to 20 libraries per lane on either Illumina HiSeq 2500 or 4000 machines, generating either 2×125bp or 2×150bp reads, respectively. Library preparation and sequencing was done at the University of North Carolina (UNC) School of Medicine’s high-throughput sequencing facility.

Raw sequence data was initially parsed and barcodes were removed by the UNC High-throughput sequencing facility. We then mapped parsed reads to each individual’s respective reference genome using the `BWA` *mem* algorithm (v0.7.15). We sorted and filtered mapped reads using `SAMTOOLS` (v1.4), marked duplicates using the `PICARD` *MarkDuplicates* tool (v2.2.4), and realigned around indels using `GATK`’s *RealignerTargetCreator* and *IndelRealigner* tools (v3.8; (McKenna et al. 2010)). Processed alignment files (.bam format) were generated separately for each individual using this pipeline.

We estimated genotypes for each individual using `GATK`’s *HaplotypeCaller* tool with options “--emitRefConfidence GVCF”, “--minReadsPerAlignmentStart 4”, “--standard_min_confidence_threshold_for_calling 8.0”, and “--minPruning 4”. We then performed joint genotyping using `GATK`’s *GenotypeGVCFs* tool. We filtered SNPs using `GATK`’s *VariantFiltration* tool with option “--filterExpression “QD < 2.0 || FS > 60.0 || SOR > 3.0 || MQ < 40.0 || MQRankSum < −12.5 || ReadPosRankSum < −8.0”” and hard-filtered sites genotyped in fewer than two individuals (`VCFtools` (v0.1.15) option “--max-missing 0.5”). To facilitate comparisons across populations where we sampled different numbers of individuals, joint genotyping and filtering was carried out on randomly selected groups of four individuals (8 chromosomes) per population, except for the population of *Z. africanus* sampled from São Tomé, where we only sampled three individuals. Lastly, for each species, we masked sites in the genome if coverage was greater than twice, or less than half, the average coverage observed across all sequenced individuals of that species.

### Estimating genetic diversity

We estimated kinship, using filtered genotypes, for all within-population pairwise comparisons using the KING method (Manichaikul et al., 2010) as implemented in the `VCFtools` “--relatedness2” tool and estimated the inbreeding coefficient (*F*) for each individual based on filtered genotypes using the `VCFtools` “--het” tool. We excluded coverage-masked sites using the “--exclude-positions” filter option. Because scaffolds in our assemblies belonging to the sex chromosomes have not been identified, we restricted our analysis of inbreeding to include only females, because homozygous genotype calls for males on the X chromosome would inflate estimates of inbreeding. In total, we estimated *F* for 7 females from the invasive range and 12 females from the native range of *Z. indianus*.

Population genetic metrics of genetic diversity were computed using `VCFtools` with coverage-masked sites excluded using the “--exclude-positions” filter option. Because *π*_*SNP*_ and *S* were highly correlated in all populations (*r* > 0.963), we focus primarily on *S*: the number of sites with segregating variation within a given 5 Kb window. We summarize estimates of genetic diversity within each population as median, 5% empirical quantile, and 95% empirical quantile values.

### Comparing genetic diversity within and outside of gene annotations

We explored the effect of being located in or around genes on levels of genetic diversity by first comparing genetic diversity across all genomic windows to diversity in windows that overlapped an annotated BUSCO gene (Waterhouse et al., 2018). We include this category of distinct annotations because these genes (2799 total) have been curated as single-copy orthologs in 25 dipteran species and we were able to annotate a high percentage (minimum 90.7%, maximum 97.3%) as being present in complete single-copies in the *de novo* assemblies we generated for this study (Table S2). Comparing diversity within these “BUSCO windows” allows for less biased comparisons between species because these windows should be less affected by aspects of genome evolution such as changes in gene copy number.

To test whether genomic regions that overlapped with gene annotations differed in levels of genetic diversity compared to regions away from genes, we used generalized linear models (GLMs) with poisson distributed error (*glm*() function in R) to model the number of segregating sites (*S*) within a genomic window as a function of the position of that window relative to a gene annotation. We classified genomic windows as overlapping, adjacent to (within 5000 kb), or distant from (> 5000kb) the nearest gene annotation. We carried out this analysis separately for each population for which we generated annotations for their species’ respective genome assembly (N = 15). Our rationale for this analysis was that either selection should have a larger effect on levels of genetic diversity in regions of the genome that contain a gene than regions that are not near a gene. For populations of *Z. indianus*, we were also interested in whether aspects of biological invasion had a different effect on levels of genetic diversity depending on the proximity of a genomic region to a gene. We therefore used a GLM to test the interaction between gene region type (i.e. overlapping, adjacent, or distant) and invasion status (i.e. invasive population of *Z. indianus*, native population of *Z. indianus*, or population of non-invasive species of *Zaprionus*) on median levels of genetic diversity across genomic windows.

### Estimating recombination rate and its effect on genetic diversity

To estimate fine-scale population recombination rates (ρ_rec_ = 2N*r*) across the genome, we used the maximum likelihood method implemented in `LDhelmet` (v1.10; (Chan et al., 2012)). For analyses involving recombination rate, we focused on populations of *Z. indianus* because we had the largest sample size of this species from a single region (Senegal: N = 14 individuals [28 haplotypes]), which allowed us to use haplotype information to estimate recombination rates across the genome (SI section 1.3). All analyses below are conducted on mean recombination rates within 5,000 bp genomic windows that were generated from median posterior estimates provided by `LDhelmet`.

We first tested for a relationship between recombination rate and gene density across 5kb windows on the 40 largest scaffolds of the *Z. indianus* genome assembly (12,080 windows) using Spearman’s rank correlation tests. We found no evidence for a relationship between recombination rate and gene density -either across all windows (Spearman’s *ρ* = −0.014; *P* = 0.12) or windows containing at least one gene (N = 4,406 windows; Spearman’s *ρ* = 0.023; *P* = 0.12). We therefore conducted independent tests for relationships between genetic diversity and recombination rate and genetic diversity within and around gene annotations.

We tested for a correlation between genetic diversity (*S*) and recombination rate, for each population of *Z. indianus*, by calculating Spearman’s ρ using the *cor*.*test* function in R. To test whether the reduction in genetic diversity we observed in invasive, relative to native, populations of *Z. indianus* varied across regions of the genome that experience different recombination rates, we calculated the correlation between median recombination rate the mean difference in S between invasive and native populations of *Z. indianus* across genomic windows. Because recombination rate was positively correlated with diversity and the number of SNPs in a genomic window affects the magnitude of change in genetic diversity that is possible, we restricted this analysis to windows with a mean number of SNPs in the across populations of *Z. indianus* in their native range between 150 and 300. To further account for differences in diversity across windows as a function of local recombination rate we scaled the observed difference in mean *S* between the invasive and native ranges of *Z. indianus* by the mean number of SNPs within genomic windows binned into five recombination rate quantiles. Our rationale for this approach was to test whether the proportional reduction in diversity in invasive populations of *Z. indianus* was greater for regions of the genome with low recombination rates, as would be expected if selection was acting to drive the genetic diversity lower in invasive populations.

### Measuring the correlation in genetic diversity between species

To facilitate interspecific comparisons, we analyzed genomic regions that spanned the first and last exon of an annotated BUSCO gene. As described above, BUSCO genes have been annotated as conserved single-copy orthologs across phylogenetically diverse Dipteran species. Focusing on BUSCO genes therefore allowed us to make direct comparisons across different species using orthologous gene regions. In total, we quantified the correlation in diversity (or lack thereof) across 2,714 genomic windows spanning orthologous BUSCO regions (hereafter “BUSCO windows”) annotated across all seven species’ genome assemblies. Correlation coefficients were estimated using the *rcorr* function from the `Hmisc` R package. To test whether comparing BUSCO windows produced correlations greater than expected by chance, we performed randomization tests where we randomly selected 2,714 genomic windows (100 iterations) and computed all pairwise Spearman’s rank correlation coefficients for those randomized windows across species. We tested whether the correlation coefficients we observed across BUSCO windows for pairwise species comparisons were outside the distribution of randomized correlation coefficients.

The genomic landscape of diversity may diverge between species with increasing phylogenetic distance due to aspects of genome evolution. To quantify whether this would affect interspecific comparisons of genetic diversity, we computed phylogenetic distances between each species using a concatenated alignment of 1709 BUSCOs shared across all species genome assemblies and representing 4,616,644 sites. Genetic distances were computed from the identity matrix of the aligned BUSCOs using the *dist*.*alignment()* function from the `seqinr` R library. We then tested whether the correlation in *S* across the genome of two species was related to their genetic distance (Spearman’s Mantel test; *mantel()* function in the `ecodist` R library).

We also calculated Spearman’s rank correlations for *S* and Tajima’s *D* between populations of *Z. indianus* (all pairwise comparisons) using diversity estimates across all 5 kb genomic window.

### Estimating thermal performance curves

To quantify differences in performance across populations, we fit a model that jointly describes the thermal performance curve to data from all populations using a hierarchical Bayesian framework (Tittes et al., 2019). This model provides estimates for the maximum height of a population’s performance curve, two shape parameters allowing for asymmetry in the performance curve, and the minimum and maximum temperatures where performance falls to 0 (T_min_ and T_max_ parameters, respectively). The height (i.e. stretch as defined in Tittes et al. 2019) reflects the maximal fitness/performance of a population. We also derived three additional parameters from the five model-estimated parameters to summarize aspects of fitness and the thermal niche: the temperature at which a population displays maximal performance (T_optimum_), the total area under the estimated performance curve (*A*_*c*_), and the breadth of the curve between the lower 25% and the upper 25% of the curve (*B*_*50*_). We estimated performance curves from data on the total number of offspring produced by a single pair of flies across mean hourly temperatures ranging from 13.9 to 28.9°C (SI). We compared performance curves between species by generating posterior draws of parameter estimates. Two populations were considered to differ with respect to parameters describing their thermal performance curve when a given parameter estimate had a probability greater than 0.95 of being different between the two populations.

## Acknowledgements

We thank M. Cenzer, C. Maxwell, A. Serrato-Capuchina, S. Yeap, E. Behrman, P. Schmidt, B. Cooper, K. Deitz, J. Coughlan, and M. Schumer for scientific discussions that improved this manuscript and/or for help in the field. This work was funded by an NSF Dimensions of Biodiversity award (1737752) to DRM and AHH. The funders had no role in an aspect of study design, data collection and analysis, or decisions with respect to publication.

## Competing interests

None of the authors declare any competing interests, financial or otherwise.

