## Supplementary Information for "Genetic diversity and thermal performance in invasive and native populations of African fig flies"

### 1. Supplementary Methods

#### *1.1 Sampling procedure*

Live flies were aspirated directly from traps and within one hour of collection, anesthetised with flynap (triethylamine in alcohols; Carolina Biological Supply) and identified under light microscopes. Up to 50 individual females from the genus *Zaprionus* (per sample location) were moved into vials containing hydrated instant drosophila media to establish isofemale lines. Males and excess females were preserved in 100% ethanol.

#### *1.2 Genome assembly*

For each species, we extracted genomic DNA from pools of 5 to 30 male flies from a single isofemale line and sequenced those extractions using Illumina and Oxford Nanopore (ONT) sequencers or, for a subset of species, ONT sequencers only (Table S1). For species with both Illumina and ONT data, we generated initial assemblies with SPAdes v3.12.0 (Bankevich et al., 2012). Assembled contigs were processed by Redundans v0.13c (Pryszcz and Gabaldón, 2016) to remove residual redundant haplotypes. Nanopore reads were corrected using FMLRC (Wang et al., 2018) and used to scaffold assembled contigs using LINKS v1.8.5 (Warren et al., 2015) with the recommended iterative approach (https://github.com/bcgsc/LINKS). Resulting scaffolds were corrected, and consensus sequences were generated, using Racon v1.3.2 (Vaser, Sović, Nagarajan, & Šikić, 2017) and Pilon v2.11 (Walker et al., 2014). For species with nanopore-only data, we mapped reads in a pairwise fashion using Minimap2 v2.15-r905 (Li, 2018, p. 2) and assembled with Miniasm v0.3-r179 (Li, 2016). We then corrected and generated consensus genome sequences with four iterations of Racon v1.3.2 (Vaser et al., 2017) followed by Medaka v0.6.2 (https://github.com/nanoporetech/medaka), respectively.

#### *1.3 Annotation*

We isolated total RNA using a standard TRIzol protocol from sex-specific groups of one day old adult flies (2 to 5 flies per extraction) that were flash frozen in liquid nitrogen. Stranded RNA-seq libraries were then constructed for each pool and sequencing was carried out on two lanes of an Illumina HiSeq 2500, run in rapid mode with 2 x 150 cycles. This sequencing approach generated between 21 and 32 million reads per extraction. Library construction and sequencing was carried out at the University of North Carolina Medical School's High-Throughput Sequencing Facility. We assembled a transcriptome for *Z. africanus* (1m, 2f pools), *Z. tuberculatus* (2f pools), *Z. nigranus* (2m, 2f pools), *Z. indianus* (2m, 2f pools), and *Z. tsacasi* (1m, 1f pools) using Trinity (Grabherr et al., 2011; Haas et al., 2013) run with default parameters. We then used MAKER (v3.01.02; (Campbell et al., 2014; Holt and Yandell, 2011)) to annotate each genome, including RepeatMasker (v4.07; (Smit and Green, 2013)) and est2genome to directly predict genes from assembled transcripts. Functional annotations were predicted using BLASTP (v2.7.1) against Swiss-Prot with an e-value cutoff of 0.000001.

#### *1.4 Estimating recombination rates across the* Z. indianus *genome*

We estimated population recombination rates ($\rho_{rec} = 2Nr$) across the *Z. indianus* genome using the maximum likelihood method implemented in LDhelmet (v1.10; (Chan et al., 2012)). Before running LDhelmet, we generated phased haplotypes for the 14 *Z. indianus* sampled from Senegal (generating 28 phased haplotypes) using read-aware phasing (`Shapeit v2.837`; (Delaneau et al. 2013)). Phasing was carried out for the 40 largest scaffolds of the *Z. indianus* assembly, totalling 61.2 Mb of sequence or ~42% of the genome. We then ran LDhelmet on the phased data by first generating haplotype configuration files for each individual using the "find_confs" script, specifying a window size of 50 SNPs. We then computed lookup tables using the "table_gen" script, specifying a population-scaled mutation rate of $\theta = 0.034$ (Watterson's $\theta$; estimated from the data using ANGSD), and a grid of recombination rate values of [0.0 0.1 10.0 1.0 100.0]. We estimated Padé coefficients, specifying $\theta = 0.034$ and 11 replicates. Finally, we estimated recombination rates, running LDhelmet's rjMCMC algorithm with a block penalty of 10 and a burn-in of 100,000 MCMC iterations followed by 1,000,000 MCMC iterations and extracted the mean recombination rate estimate between each pair of SNPs using LDhelmet's "post_to_text" script. To summarize variation in recombination rate across the genome, we first removed unrealistically high estimates of $\rho_{rec}$ (i.e. $\rho_{rec} > 1$, corresponding to a recombination rate greater than ~ $15*10^{-8}$) and then calculated the mean recombination rate in 5000 bp windows across the genome from median estimates provided by LDhelmet.

*1.5 Measuring thermal performance*

We measured adult-to-adult performance under four different temperatures and a 10:14 hour night:day light cycle. Temperatures were (night:day) 11°C:16°C, 16°C:21°C, 21°C:26°C, and 26°C:31°C, resulting in mean hourly temperatures of 13.9, 18.9, 23.9, and 28.9°C, respectively. To initiate the experiment, individuals were collected as one to three day old adults, briefly anesthetized with $CO_2$, and placed as individual pairs into vials containing standard cornmeal agar medium. Each pair was then allowed to recover from anesthesia for 12 to 24 hours at room temperature before being randomly assigned to one of the four temperature treatments. Each pair of flies was then allowed to lay eggs for 7 to 9 days, after which they were removed from the vials and a dampened kimwipe was added to each vial as a pupation site for the larvae. We then counted the total number of offspring that successfully enclosed within each vial. We measured adult-to-adult performance in this way for a total of 900 pairs, with an average of 9 pairs per temperature per species. Temperature and light was controlled using Percival incubators (model DR-36VL). Relative humidity within the incubators was negatively correlated with temperature, but was maintained between 80% and 50%.

We modified the original model presented in (Tittes et al., 2019) to account for higher variability in reproduction and survival in our dataset compared to the data on which the model was originally developed. We modeled the data as a mixture of a Gaussian probability density that described thermal performance and a Bernoulli probability mass that described excess zeros caused by mortality and failure of pairs to reproduce, which we will subsequently refer to as "mortality" for simplicity. We assumed the probability of mortality was inversely proportional to the mean thermal performance, such that more zeros are expected to occur near the thermal tolerance limits. The Bayesian p-value for the model was 0.77, indicating an adequate goodness-of-fit of the model to the data. Other modeling details including values chosen for priors remained the same as in Tittes et al. (2019). We have posted the Stan code that provides a precise description of the model at: https://github.com/silastittes/performr/tree/zin.

In the main text, we report on parameter estimates of thermal performance minimum ($T_{min}$ == x_min), thermal maximum ($T_{max}$ == x_max), thermal optimum ($T_{optimum}$ == maxima), maximum realized fitness (max. fitness == stretch), and a measure of thermal niche breadth ($B_{50}$), each derived from the estimated thermal performance curves. Each of these parameters, except for $B_{50}$, is described in Tittes et al. (2019). We estimated $B_{50}$ as the difference in temperature values that captured the central 50% of the curve area, and was calculated as the difference between
critical = [(1 - ((1 - X_CRITICAL) ^ (1/shape_2)))^(1/shape_1)] * (x_max - x_min) + x_min,
where X_CRITICAL was chosen to be 0.75 0.25, respectively.

**Table S1.** Summary of the data used to generate each of the seven draft genome assemblies reported in the main text. The amount of sequence data generated for each species and each sequencing technology is given in billions of base pairs (Gbp). For nanopore sequencing we also report the number of reads in millions (M) and the mean read length in base pairs.

| species | Collection location | Illumina read type | Illumina amount | Nanopore reads (N) | Nanopore amount | Nanopore mean read length |
|---|---|---|---|---|---|---|
| *Z. indianus* | Florida | MiSeq 250 x 2 | 8.6 Gbp | 1.7M | 4.2 Gbp | 2471 |
| *Z. africanus* | Sao Tome | n/a | n/a | 1.7M | 7.4 Gbp | 4259 |
| *Z. nigranus* | Sao Tome | MiSeq 300 x 2 | 11.1 Gbp | 1.6M | 1.9 Gbp | 1250 |
| *Z. taronus* | Sao Tome | n/a | n/a | 2.5M | 10.0 Gbp | 3900 |
| *Z. inermis* | Sao Tome | n/a | n/a | 3.1M | 10.6 Gbp | 3400 |
| *Z. tuberculatus** | Zambia | n/a | n/a | n/a | n/a | n/a |
| *Z. tsacasi* | Sao Tome | HiSeq 2500 150 x 2 | 14.9 Gbp | .945M | 2.5 Gbp | 2646 |

* The *Z. tuberculatus* assembly was generated by Dovetail genomics with their proprietary Chicago libraries, a Hi-C library, and Illumina sequence data.

**Table S2.** Summary statistics for each of the seven draft genome assemblies reported in the main text. BUSCO annotation report is based on a total of 2799 single copy orthologs curated in 25 genomes of different species of Diptera. Annoations were generated using RNA-seq data, Trinity, and the MAKER annotation pipeline (see Supplementary Methods and the Main Text).

| species | Assembly size (Mbp) | # contigs | N50 (bp) | % complete BUSCO | % complete single-copy BUSCO | % duplicated BUSCO | # annotated transcripts |
|---|---|---|---|---|---|---|---|
| *Z. indianus* | 145.7 | 649 | 773,890 | 96.6 | 96.1 | 0.5 | 10,013 |
| *Z. africanus* | 167.6 | 689 | 1,499,604 | 95 | 94.1 | 0.9 | 10,424 |
| *Z. nigranus* | 142.9 | 2553 | 776,169 | 97.8 | 97.3 | 0.5 | 9,769 |
| *Z. taronus* | 187.9 | 536 | 2,214,536 | 95.2 | 93.7 | 1.5 | 9,275 |
| *Z. inermis* | 165.5 | 572 | 2,453,702 | 94.9 | 94 | 0.9 | n/a |
| *Z. tuberculatus* | 176.2 | 880 | 25,350,852 | 93.2 | 90.7 | 2.5 | 11,071 |
| *Z. tsacasi* | 150.1 | 1269 | 335,836 | 96.3 | 95.7 | 0.6 | 10,408 |

**Table S3.** Populations and sample sizes for which whole genome resequencing data was generated to estimate genetic diversity. Sample sizes are reported as the number of chromosomes sampled from each population (i.e. 2 x the number of individuals sampled).

| species | location | 2N |
|---|---|---|
| *Z. indianus* | Hawaii | 8 |
| *Z. indianus* | North Carolina | 12 |
| *Z. indianus* | Tennessee | 8 |
| *Z. indianus* | Sao Tome | 12 |
| *Z. indianus* | Senegal (forest) | 14 |
| *Z. indianus* | Senegal (desert) | 14 |
| *Z. indianus* | Kenya | 14 |
| *Z. indianus* | Zambia | 12 |
| *Z. africanus* | Sao Tome | 6 |
| *Z. africanus* | Kenya | 10 |
| *Z. nigranus* | Sao Tome | 10 |
| *Z. taronus* | Sao Tome | 22 |
| *Z. inermis* | Sao Tome | 8 |
| *Z. tuberculatus* | Senegal | 14 |
| *Z. tuberculatus* | Sao Tome | 14 |
| *Z. tsacasi* | Sao Tome | 8 |

**Table S4.** Summary of genetic diversity within each population. Population identifiers are in the format species-POPULATION-replicate, where species is abbreviated as "z" followed by the first three letters of the species name and population is the abbreviation of the collection location. Summary statistics were calculated from groups of four individuals and in populations where more than four individuals were sampled, multiple subsamples (replicates) were analyzed. Mean, 5% empirical quantile (5%), and 95% empirical quantile (95%) for nucleotide diversity (*pi*), the number of segregating sites (*S*), and Tajima's *D* (T. *D*) calculated in 5 kb windows across the genome are reported.

| population | *pi* median | *pi* (5%) | *pi* (95%) | *S* median | *S* (5%) | *S* (95%) | T. *D* median | T. *D* (5%) | T. *D* (95%) |
|---|---|---|---|---|---|---|---|---|---|
| zafr-KEN-1 | 0.0169 | 0.0011 | 0.0276 | 231 | 3 | 366.2 | -0.66 | -1.10 | 0.11 |
| zafr-KEN-2 | 0.0173 | 0.0011 | 0.0279 | 235 | 4 | 371 | -0.68 | -1.13 | 0.09 |
| zafr-ST-1 | 0.0188 | 0.0010 | 0.0304 | 205 | 3 | 332 | -0.42 | -0.93 | 0.49 |
| zind-HI-1 | 0.0110 | 0.0006 | 0.0206 | 120 | 0 | 231 | 0.51 | -1.28 | 2.09 |
| zind-KEN-1 | 0.0171 | 0.0030 | 0.0274 | 219 | 37 | 339 | -0.54 | -1.05 | 0.17 |
| zind-KEN-2 | 0.0175 | 0.0029 | 0.0279 | 230 | 36 | 349 | -0.67 | -1.15 | 0.02 |
| zind-NC-1 | 0.0138 | 0.0016 | 0.0241 | 160 | 10 | 281 | 0.17 | -1.02 | 1.71 |
| zind-NC-2 | 0.0137 | 0.0014 | 0.0241 | 159 | 7 | 281 | 0.14 | -1.18 | 1.64 |
| zind-SEN desert-1 | 0.0174 | 0.0021 | 0.0275 | 230 | 28 | 347 | -0.71 | -1.18 | -0.17 |
| zind-SEN desert-2 | 0.0175 | 0.0022 | 0.0276 | 231 | 28 | 348 | -0.70 | -1.17 | -0.14 |
| zind-SEN forest-1 | 0.0176 | 0.0021 | 0.0278 | 232 | 28 | 350 | -0.71 | -1.18 | -0.17 |
| zind-SEN forest-2 | 0.0176 | 0.0021 | 0.0279 | 233 | 28 | 350 | -0.72 | -1.18 | -0.15 |
| zind-ST-1 | 0.0161 | 0.0018 | 0.0271 | 206 | 21 | 336 | -0.48 | -1.05 | 0.58 |
| zind-ST-2 | 0.0161 | 0.0018 | 0.0271 | 206 | 21 | 335 | -0.48 | -1.04 | 0.54 |
| zind-TN-1 | 0.0136 | 0.0015 | 0.0241 | 156 | 4 | 280.65 | 0.22 | -0.92 | 1.93 |
| zind-ZAM-1 | 0.0168 | 0.0026 | 0.0276 | 222 | 32 | 349 | -0.66 | -1.16 | 0.45 |
| zind-ZAM-2 | 0.0172 | 0.0027 | 0.0278 | 229 | 33 | 350 | -0.71 | -1.22 | -0.12 |
| zine-ST-1 | 0.0031 | 0.0001 | 0.0084 | 35 | 0 | 92 | 0.52 | -1.74 | 2.07 |
| znig-ST-1 | 0.0018 | 0.0005 | 0.0045 | 23 | 6 | 55 | 0.16 | -0.90 | 1.29 |
| znig-ST-2 | 0.0019 | 0.0005 | 0.0045 | 23 | 6 | 55 | 0.12 | -0.92 | 1.14 |
| ztar-ST-1 | 0.0095 | 0.0003 | 0.0219 | 111 | 0 | 260 | -0.34 | -1.17 | 0.42 |
| ztar-ST-2 | 0.0092 | 0.0003 | 0.0216 | 107 | 0 | 257 | -0.30 | -1.15 | 0.43 |

| | | | | | | | | | |
|---|---|---|---|---|---|---|---|---|---|
| ztar-ST-3 | 0.0090 | 0.0003 | 0.0216 | 100 | 0 | 256 | -0.20 | -1.06 | 0.79 |
| ztsa-ST-1 | 0.0115 | 0.0007 | 0.0214 | 143 | 5 | 257 | -0.21 | -0.88 | 0.53 |
| ztub-SEN-1 | 0.0100 | 0.0003 | 0.0191 | 133 | 2 | 247 | -0.41 | -1.03 | 0.42 |
| ztub-SEN-2 | 0.0093 | 0.0003 | 0.0179 | 126 | 1 | 239 | -0.33 | -0.89 | 0.61 |
| ztub-ST-1 | 0.0122 | 0.0003 | 0.0223 | 152 | 1 | 272 | -0.37 | -0.89 | 0.52 |
| ztub-ST-2 | 0.0122 | 0.0003 | 0.0224 | 152 | 2 | 273 | -0.39 | -0.97 | 0.39 |

**Table S5.** Summary of genetic diversity within each population, as reported in Table S4, but restricted to genomic windows overlapping an annotated BUSCO gene. Only one replicate of four randomly selected individuals was run for each population, as results in Table S4 indicate that there was not a large variance between estimates generated from different subsamples of four individuals.

| population | *pi* (median) | *pi* (5%) | *pi* (95%) | *S* (median) | *S* (5%) | *S* (95%) | T. *D* (median) | T. *D* (5%) | T. *D* (95%) |
|---|---|---|---|---|---|---|---|---|---|
| zind-HI | 0.0099 | 0.0018 | 0.0203 | 109 | 0 | 231 | 0.48 | -1.26 | 2.10 |
| zind-NC | 0.0123 | 0.0031 | 0.0241 | 147 | 28.85 | 281.15 | 0.15 | -0.98 | 1.69 |
| zind-TN | 0.0121 | 0.0029 | 0.0241 | 142 | 20 | 281 | 0.24 | -0.77 | 1.96 |
| zind-ST | 0.0147 | 0.0032 | 0.0276 | 193 | 43 | 342 | -0.51 | -1.09 | 0.36 |
| zind-SEN desert | 0.0156 | 0.0052 | 0.0278 | 213 | 79 | 351 | -0.72 | -1.21 | -0.28 |
| zind-SEN forest | 0.0157 | 0.0052 | 0.0282 | 215 | 80 | 356 | -0.72 | -1.21 | -0.28 |
| zind-KEN | 0.0153 | 0.0053 | 0.0275 | 203 | 74.7 | 341.15 | -0.55 | -1.06 | 0.11 |
| zind-ZAM | 0.0152 | 0.0051 | 0.0280 | 206 | 71.85 | 353.15 | -0.67 | -1.22 | 0.15 |
| zafr-ST | 0.0196 | 0.0056 | 0.0319 | 221 | 67 | 354 | -0.49 | -0.90 | 0.11 |
| zafr-KEN | 0.0178 | 0.0050 | 0.0296 | 251 | 81 | 394.3 | -0.71 | -1.13 | -0.26 |
| ztub-ST | 0.0124 | 0.0046 | 0.0211 | 160 | 61 | 263 | -0.40 | -0.84 | 0.14 |
| ztub-SEN | 0.0109 | 0.0037 | 0.0192 | 150 | 50 | 252 | -0.45 | -0.87 | 0.23 |
| zinerm-ST | 0.0029 | 0.0001 | 0.0070 | 33 | 1 | 77 | 0.53 | -1.80 | 2.08 |
| ztsac-ST | 0.0095 | 0.0017 | 0.0206 | 121 | 24 | 247 | -0.21 | -0.97 | 0.39 |
| znig-ST | 0.0013 | 0.0004 | 0.0036 | 17 | 4 | 44 | 0.09 | -1.03 | 1.19 |
| ztar-ST | 0.0064 | 0.0011 | 0.0194 | 85 | 17 | 237 | -0.36 | -1.29 | 0.35 |

**Table S6**. Mean number of segregating sites (*S*) across 5 kb genomic windows grouped by their location relative to an annotated gene. Windows were classified as overlapping a gene, within 5 kb, but not overlapping (adjacent), or further than 5kb from a gene (distant). The last two columns show relative amounts of genetic diversity contained within windows that overlapped an annotated gene and either adjacent or distant genomic windows.

| population | overlapping | adjacent | distant | overlapping/adjacent | overlapping/distant |
|---|---|---|---|---|---|
| zind-HI | 129 | 134 | 133 | 0.9626866 | 0.9699248 |
| zind-NC | 168 | 176 | 172 | 0.9545455 | 0.9767442 |
| zind-TN | 166 | 172 | 167 | 0.9651163 | 0.994012 |
| zind-ST | 221 | 230 | 224 | 0.9608696 | 0.9866071 |
| zind-SENdesert | 246 | 257 | 256 | 0.9571984 | 0.9609375 |
| zind-SENforest | 249 | 260 | 258 | 0.9576923 | 0.9651163 |
| zind-KEN | 233 | 242 | 243 | 0.9628099 | 0.9588477 |
| zind-ZAM | 238 | 248 | 243 | 0.9596774 | 0.9794239 |
| zafr-ST | 237 | 230 | 214 | 1.0304348 | 1.1074766 |
| zafr-KEN | 274 | 264 | 248 | 1.0378788 | 1.1048387 |
| ztub-ST | 161 | 167 | 173 | 0.9640719 | 0.9306358 |
| ztub-SEN | 146 | 145 | 145 | 1.0068966 | 1.0068966 |
| ztsac-ST | 138 | 153 | 168 | 0.9019608 | 0.8214286 |
| znig-ST | 20 | 24 | 27 | 0.8333333 | 0.7407407 |
| ztar-ST | 102 | 130 | 152 | 0.7846154 | 0.6710526 |

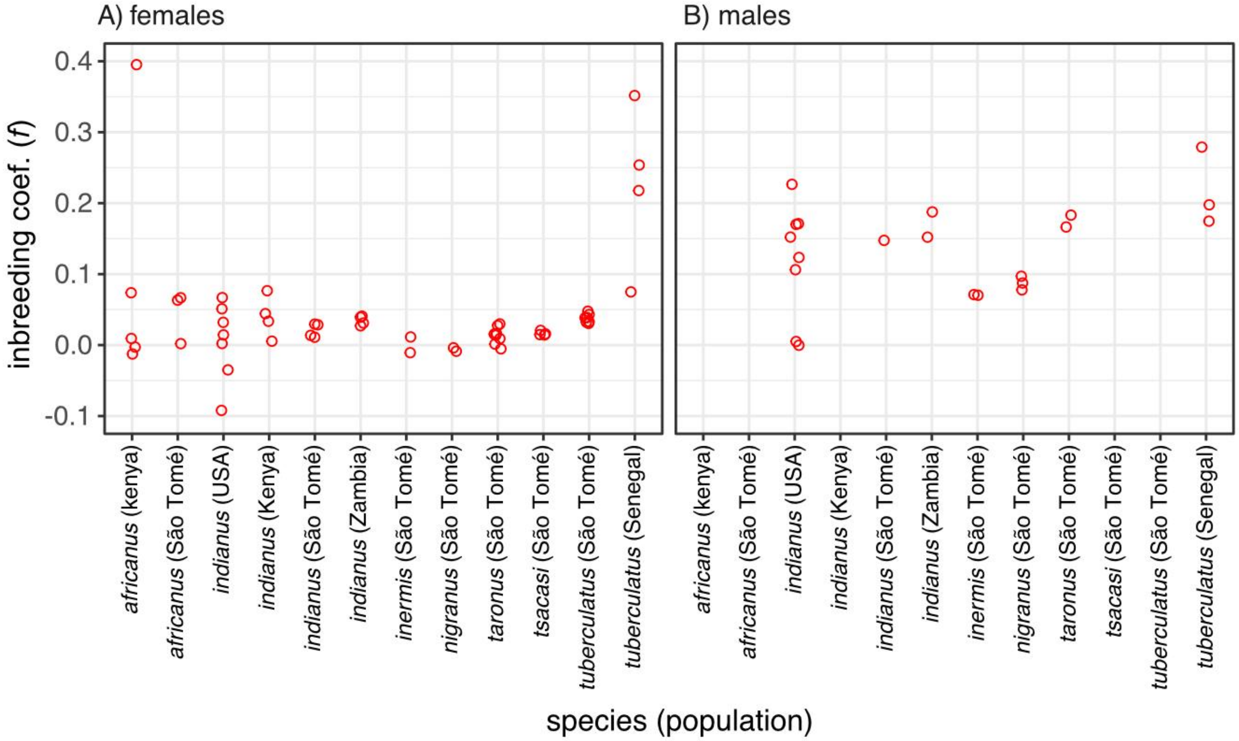


**Figure S1.** Estimated inbreeding coefficient (*f*) for all sequenced individuals with known sex at time of sequencing (A: females; B: males). Inbreeding coefficients were estimated using the KING method as implemented in VCFTools (see Main Text for details).

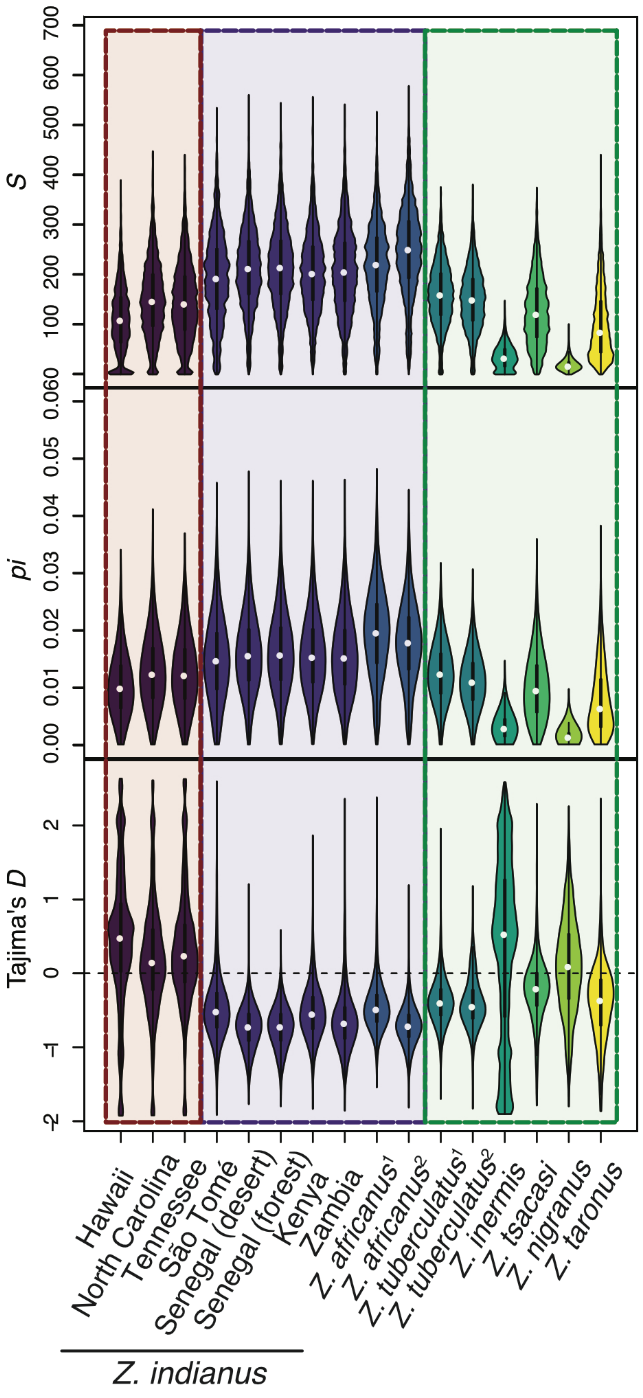


**Figure S2.** Estimates of genetic diversity summarized across 5 kb genomic windows that overlap with an annotated BUSCO. Shaded boxes group populations as invasive *Z. indianus* (three leftmost violins), native *Z. indianus* and *Z. africanus* (seven central violins), and other species (six rightmost violins). See Figure 2 in the main text for results across the entire genome.

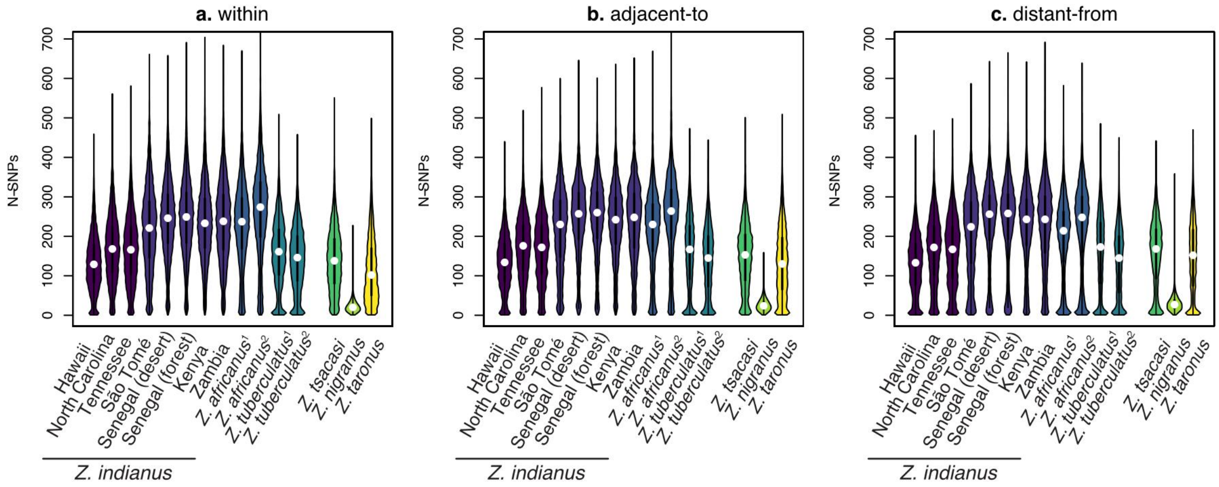


**Figure S3**. Genetic diversity (*S* == N-SNPs) summarized for windows either overlapping an annotated gene (**a**), within 5kb of an annotated gene (**b**), or greater than 5kb from the nearest annotated gene (**c**).

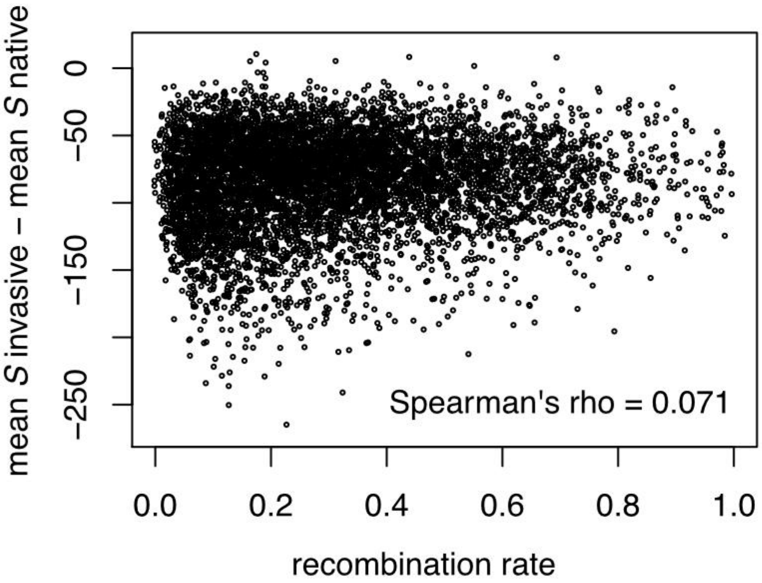


**Figure S4.** Correlation between the difference in amounts of genetic diversity (the number of segregating sites: *S*; mean across invasive populations - mean across native populations) and (population) recombination rate for *Z. indianus*.

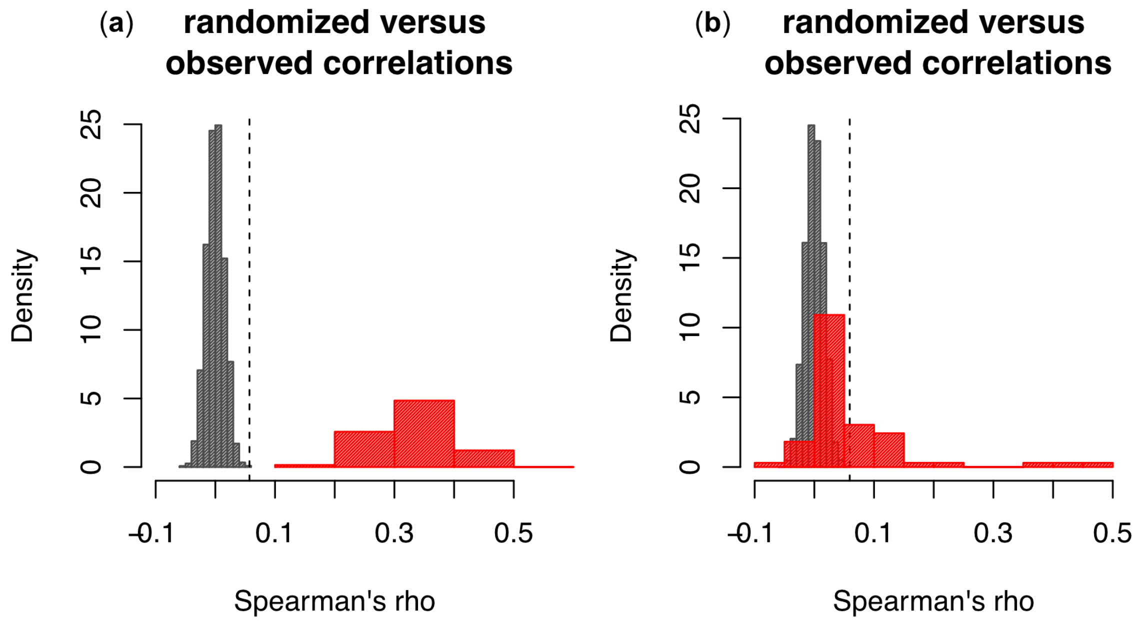


**Figure S5**. Randomized and observed correlation coefficients for genetic diversity (S; panel (a)) and Tajima's *D* (b). Grey histograms in each panel show the distribution of correlation coefficients generated when randomly selecting genomic windows that span BUSCO annotated genes in two species' genomes. Red histograms show observed correlation coefficients across all pairwise interspecific comparisons. The dashed vertical line in each panel represents the 95% tail of the randomized distribution.
